# Discovering cell types and states from reference atlases with heterogeneous single-cell ATAC-seq features

**DOI:** 10.1101/2025.09.19.677191

**Authors:** Yuqi Cheng, Xiuwei Zhang

## Abstract

Despite substantial recent advances in query mapping and cell type or cell state discovery tools, their application to single-cell assay for transposase-accessible chromatin using sequencing (scATAC-seq) data remains challenging. The heterogeneous nature of peak feature spaces across samples hinders the effectiveness of existing methods, while the absence of dedicated tools for detecting perturbed cell types and states in scATAC-seq data further limits the depth of downstream analyses. To address these limitations, we present EpiPack, an integrative computational toolkit that leverages heterogeneous transfer learning and graph-based modeling strategies to advance scATAC-seq analysis. At its core, the Peak Embedding Informed Variational Inference (PEIVI) framework within EpiPack enhances mappable reference construction, query mapping, and label transfer, demonstrating that leveraging heterogeneous features in scATAC-seq data outperforms methods relying solely on conventional homogeneous features. In addition, EpiPack’s global–local out-of-reference (OOR) detection framework achieves robust and efficient detection of perturbed cell types and states, extending the utility of scATAC-seq to disease and perturbation contexts. With its modular design and transferable pre-trained references, EpiPack can be readily applied to diverse analytical tasks and is available as a Python package at https://github.com/ZhangLabGT/EpiPack.

## Main

Recent advancements in scATAC-seq technology, by revealing regulatory elements^1–3^and biological networks^4, 5^at single-cell resolution, have revolutionized epigenetic research. To learn biological insights from a scATAC-seq dataset, discovering cell types and states is the initial key step. Reference mapping methods, where a newly generated dataset to be analyzed, called the query data, is mapped onto curated large-scale reference data, have the advantage of automated cell type annotation without the need of manual steps or biomarkers provided. Meanwhile, perturbed cell clusters and states in the query data can also be distinguished from the reference control. With the rapid generation of large-scale systematic scATAC-seq cell atlases^3, 6, 7^, the direct discovery and transfer of cell types from scATAC reference data has become increasingly imperative. However, while similar efforts for such tasks have been extensively developed in single-cell RNA sequencing (scRNA-seq) data^8–13^, learning cell types and states from massive-scale reference atlases still face significant challenges in scATAC-seq data.

Computationally, the divergences in feature spaces across different scATAC-seq datasets pose a major obstacle to the assembly of reference atlases and the mapping of query data. This is not an issue in the case of scRNA-seq data since all scRNA-seq features are from the same gene list for the same species. But the feature space of a scATAC-seq dataset is composed of various accessibility peaks (genomic regions) that are specific to this dataset; we cannot directly obtain a consistent feature set intersection across more than one scATAC-seq dataset. However, current mapping tools and classifiers rely entirely on obtaining homogeneous aligned feature spaces^14^, forcing the scATAC-seq data to compromise information enrichment for feature transformation. For instance, the most common practice is using gene activity score^15–17^. While this strategy makes feature alignment straightforward and thus facilitates label transfer to new query data, it has been shown to lead to significant information loss^18^. An alternative is to use merged overlapping peaks across samples^19–21^. While this approach aims to use peak-level data matrices, which are more informative than transforming peak information into gene activity scores, a few challenges still remain. First, obtaining shared peak features and building the pretrained reference model can be inefficient given the extremely high dimensions of each dataset^18^; second, given the heterogeneity of peak features across data matrices, intersecting the peak feature set of query data with the reference model may lead to insufficient overlapping features. Moreover, the query and reference datasets can even be derived from different reference genomes (e.g., Hg19 vs. Hg38), which can further reduce the amount of shared peak features. Therefore, while relying on homogeneous features to construct reference atlases and to map query data is effective and straightforward in scRNA-seq, it appears impractical for scATAC-seq. The issue of feature heterogeneity in scATAC-seq has been largely overlooked and remains underexplored.

The lack of tools capable of identifying cell types or cell states that are not measured in the reference datasets for scATAC-seq data presents another challenge. Recent methods developed for scATAC-seq cell type annotation, such as Cellcano^15^and EpiAnno^16^, are limited to labeling cells already present in the reference datasets. However, out-of-reference (OOR) cells are often considered more critical for gaining biological insights into the epigenetic mechanism^22^, particularly in disease or developmental studies. Meanwhile, although progress has been made in developing tools for novel cell type detection^11, 12, 23^and differential abundance analysis^24–26^(also known as perturbed cell state detection) in the scRNA-seq field, most of these methods are designed as end-to-end solutions tailored to gene expression data thus cannot be easily integrated into a pipeline for scATAC-seq data^27, 28^. Even methods that can do so, in principle, such as Milo^24^and MELD^26^still suffer from critical limitations. Milo is limited to coarse subpopulation-level testing, while MELD relies on a fixed kernel that risks oversmoothing and lacks significance testing and FDR control. This highlights the pressing need for an OOR detection tool integrated in a scATAC-seq annotation platform.

To address these challenges, we introduce EpiPack, a comprehensive deep-learning toolkit for scATAC-seq data reference mapping, cell-type automated annotation, and OOR cell types/states discovery without the requirement of aligned peak features. In particular, to solve the issue of disparate feature spaces between the source and target domains, EpiPack employs Peak Embedding Informed Variational Inference (PEIVI), a heterogeneous transfer learning paradigm coupled with conditional generative modeling, utilizing a bridge architecture to leverage distinct peak features between datasets and learn more informative embedding spaces. Benchmark tests demonstrated the superiority of this design over tools using homogeneous feature space for reference mapping, label transfer, and OOR detection in scATAC-seq data. Furthermore, through pseu- do-perturbation experiments, we showed that both the global and local OOR detectors in EpiPack achieve high sensitivity, robust FDR control, and reduced running time. In addition, we demonstrated the full features of EpiPack by constructing a reference model from publicly available healthy PBMC datasets and uncovering disease-associated perturbation clusters within the COVID sample mapping space. EpiPack is released as an open-source and user-friendly Python package, available at https://github.com/ZhangLabGT/EpiPack

## Results

### Overview of EpiPack

EpiPack leverages both gene activity score matrices, which enable feature bridging across datasets, and peak-level matrices, which preserve fine-resolution chromatin accessibility. It takes heterogeneous reference and query scATAC-seq datasets with distinct peak sets and their gene score matrices as inputs. The workflow of EpiPack is illustrated in Fig.1a. The *PEIVI* module integrates multi-source scATAC-seq references into a mappable space, generating a pre-trained base model. This model can be fine-tuned to map query data onto a joint embedding space, enabling label transfer via a metric learning-based *classifier*. Meanwhile, a distance-based *global OOR detector* and a graph-based *local OOR detector* are integrated for detecting novel cell types or subtle state shifts.

**Fig. 1.**
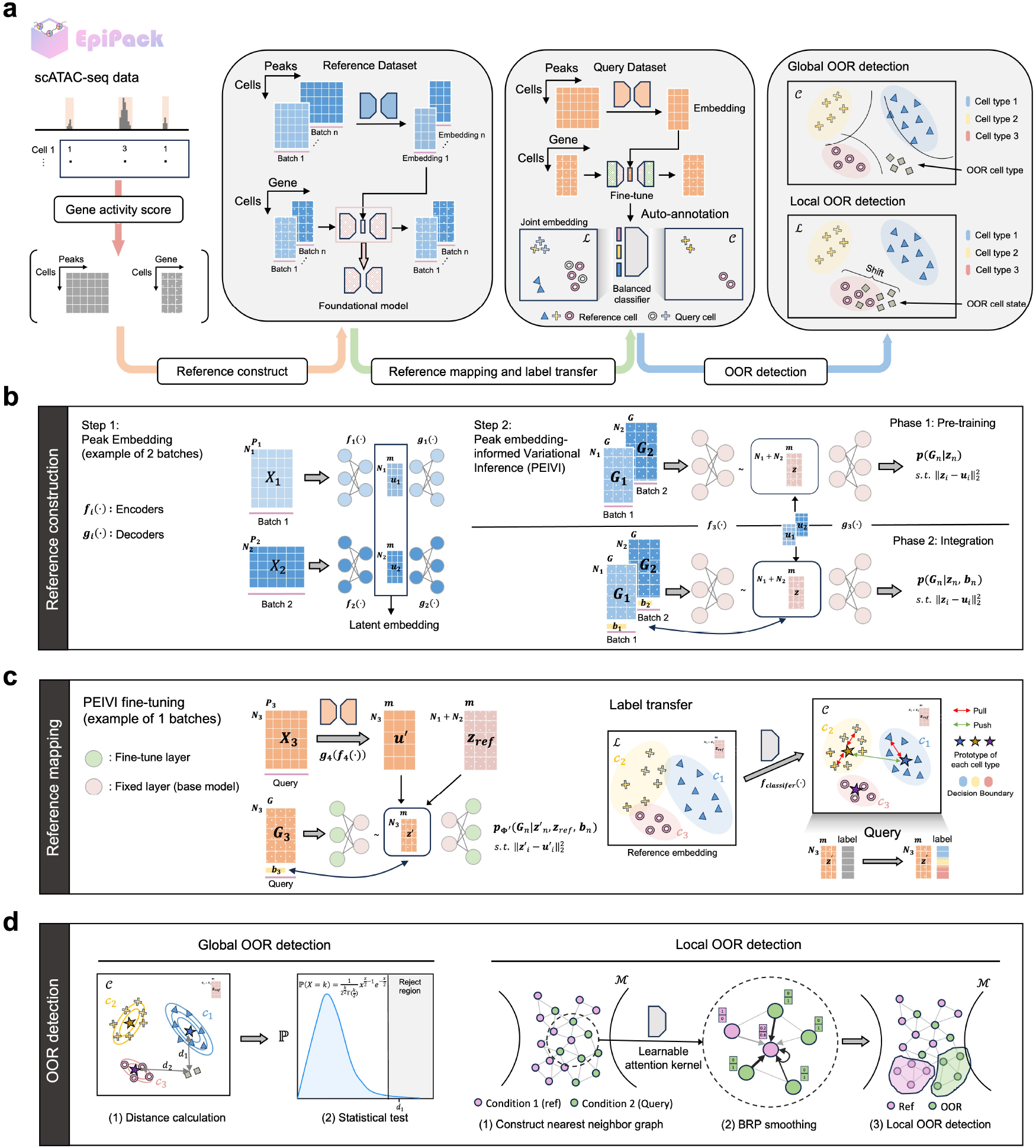
Architecture of the EpiPack framework. **a**. Workflow and functionalities of the EpiPack framework. EpiPack comprises four core algorithms: EpiPack *PEIVI* for constructing references and query mappings, EpiPack *classifier* for cell type transfer and providing a distance space, EpiPack *global OOR detector* and EpiPack *local OOR detector* for OOR detection. **b**. PEIVI integrates peak embedding information into latent factors using a conditional generative model constrained by heterogeneous information. The pretrained models and reference atlas are readily available for downstream query mapping. **c**. PEIVI completes heterogeneous transfer learning tasks by fine-tuning the pre-trained model. A supervised classifier based on metric learning provides a separable classification space by optimizing intra-class angles and inter-class distances, enabling accurate cell type annotation. **d**. The Global OOR detector is coupled with the classification space 𝒞, providing statistically significant and interpretable uncertainty scores in the global metric space using the Mahalanobis distance. In contrast, the Local OOR detector models nearest-neighbor graphs in the joint embedding space ℳ, leveraging a learned kernel and BRP smoothing techniques to compute uncertainty scores within continuous non-Euclidean manifolds and identify perturbed cell states.

#### PEIVI enables query mapping with heterogeneous features

EpiPack PEIVI aims to construct a mappable reference that can both accommodate atlas-scale query mapping and preserve peak domain information. Previous studies^15, 17, 29^showed that the gene score matrix of each scATAC-seq dataset can provide homologous and essential information that is similar to peak embeddings though at a lower resolution, which thus can be a natural and effective bridge to link the pre-generated peak embedding spaces of different reference sources. Taking advantage of this assumption, PEIVI builds the heterogeneous model based on the conditional variational autoencoder (CVAE) architecture^30, 31^. By modeling latent factors generated from the gene score space through probabilistic variational inference, PEIVI integrates the corresponding precomputed peak embeddings *u*_*i*_ as an additional prior constraint (Fig.1a). This prior is incorporated into the latent representation, ensuring that each encoded latent space is enriched with the peak-level regulatory information associated with its reference input (Methods). To maximize the integration of peak domain information, PEIVI adopts a stepwise training approach. It first performs pretraining under the sole constraint of *u*_*i*_ without incorporating the covariance factor *b*_*i*_ as posterior information. Next, during the harmonization process, the model trains *b*_*i*_ and *u*_*i*_ in a trainable manner to regress out batch information introduced by *u*_*i*_, ultimately returning a harmonized reference space embedding *z*_*i*_ for all the batches. (Fig.1b). This advancement allows PEIVI to take the unified gene score matrix as a scalable input to link batch-specific latent spaces, thus bypassing directly aligning peak features.

After training a model on multiple reference datasets, EpiPack employs a fine-tuning procedure to facilitate robust transfer learning, enabling the mapping of query data onto the reference embedding space (Fig.1c left and Methods). By fine-tuning the pre-trained foundational model, PEIVI mapping can approximate *de novo* integration of reference and query cells while only requiring minimal computational resources.

#### EpiPack classifier

Given the joint latent space of the reference and query datasets, we developed a metric learning-based neural network classifier to perform label transfer to the query dataset. This classifier projects embeddings from the joint latent space ℒ to the classification space 𝒞 (Fig.1c right and Methods). The classifier employs a loss function that incorporates both a logit-based angular constraint to maximize inter-class distance and a prototype-based distance constraint to enhance intra-class compactness (Methods), which aims to provide a more separable decision boundary in space 𝒞. To ensure the rare cell types are well represented, the classifier is equipped with a weighted sampling technique^32^targeting the cell population imbalance issue while training.

#### Global-Local OOR framework

**Global OOR** represents the scenario that unseen cell types exhibit significant differences from in-reference cell types (e.g., CD4 T cells vs. B cells). In the joint latent space, these OOR types appear as separate clusters rather than overlapping with or connecting to the in-reference cell types (Extended Fig.1a). In this case, the distances between OOR data points and other mapped cell types can be approximated using the L2 norm^30^. Then, in the classification space 𝒞 (which has enlarged space between clusters and can expose OOR cell types more easily), EpiPack detects OOR cells efficiently using Mahalanobis distance. Since the Mahalanobis distance follows a chi-square distribution^33^, EpiPack performs a cell-type-specific hypothesis test on the distance vector between each query cell and each existing cell type, with the probability measure representing the confidence score of the query cell belonging to each annotated cell population (Fig.1d Left). This enables EpiPack to deliver statistically significant annotation results while providing confidence scores, avoiding the overconfidence issue often observed in supervised models. The annotations from the classifier are further refined based on rejection thresholds derived from probability density functions or manually set, enabling the identification of OOR cell populations (Methods).

**Local OOR** represents the scenario that perturbed cell states exhibit non-significant differences from in-reference cell types (e.g., healthy monocytes vs. disease-state monocytes). In the joint latent space, these OOR cells form a continuous and smooth manifold together with the state-shifting clusters from the reference space (Extended Fig.1b), adhering to local Euclidean geometry rather than a globally L2 space. Thus in our local OOR detection module (Fig.1d right and Extended Fig.1c), a mutual kNN graph is first constructed between reference and query cells in the joint latent space. Each edge is characterized by a learnable multi-dimensional feature vector (e.g., inter-cell distance, local density difference), which is used to train a learnable attention kernel. A bi-directional residual propagation (BRP) kernel calculated from the attention kernel diffuses OOR scores across the graph while retaining a residual connection to each node’s original embedding to mitigate over-smoothing. The resulting OOR scores are converted into *p*-values and corrected using Benjamini–Hochberg FDR control to identify local OOR cells (Methods).

### Accurate query mapping and label transfer with heterogeneous scATAC-seq features

As mentioned earlier, the divergence in feature spaces between scATAC-seq data batches is the primary obstacle to integrating query data into reference atlases and transferring cell type labels. Therefore, we first evaluated whether our heterogeneous transfer learning framework can improve mapping performance on scATAC-seq data, compared to methods based on homogeneous features. To rigorously assess EpiPack’s performances in the scenario that reflects real experimental conditions, we applied our method to five curated peripheral blood mononuclear cell (PBMC) datasets assayed with three different 10x Genomics protocols: v1.1, v2, and multiome, which are top protocols benchmarked by Florian et al^34^compared to other experimental protocols (Methods). Since, in reality, query data can often be generated from sequencing platforms different from that of the reference, we employed a cross-platform benchmarking strategy. We set up three experiment groups: for each, datasets from two technologies were used as references and the other one as a query. We compared EpiPack to multiple popular unsupervised reference mapping and supervised cell type annotation methods, which are solely based on aligned features, including the gene-score-based models (scArches (scVI)^12^, Seurat^35^, Cellcano^15^, and SVM (support vector machine)) and the peak-based models (PeakVI^20^and Signac^36^). Among these baseline methods, scArches (scVI), PeakVI, Seurat v4, and Signac are commonly used for both reference mapping and label transfer, while Cellcano and SVM are top performers in the scATAC-seq annotation task^15^.

We first quantified the performance of EpiPack PEIVI on the unsupervised reference mapping task by using a suite of biological conservation and batch correction metrics proposed in the benchmarking platform scIB^18^. Detailed explanations of each metric are available in Methods. Compared with reference mapping tools that require aligned homogeneous features (either gene scores or overlapped peaks), our analysis revealed that PEIVI achieved the best performance in both biological conservation and batch correction metrics simultaneously and was consistently the top method across all experiment groups (Fig.2a and Supplementary Fig.1); uniform manifold approximation and projection (UMAP) visualization showed that cell populations were well-aligned in PEIVI, while the reference and query datasets are precisely mixed (Fig.2d and Supplementary Fig.2), which further confirmed the mapping performance of EpiPack.

**Fig. 2.**
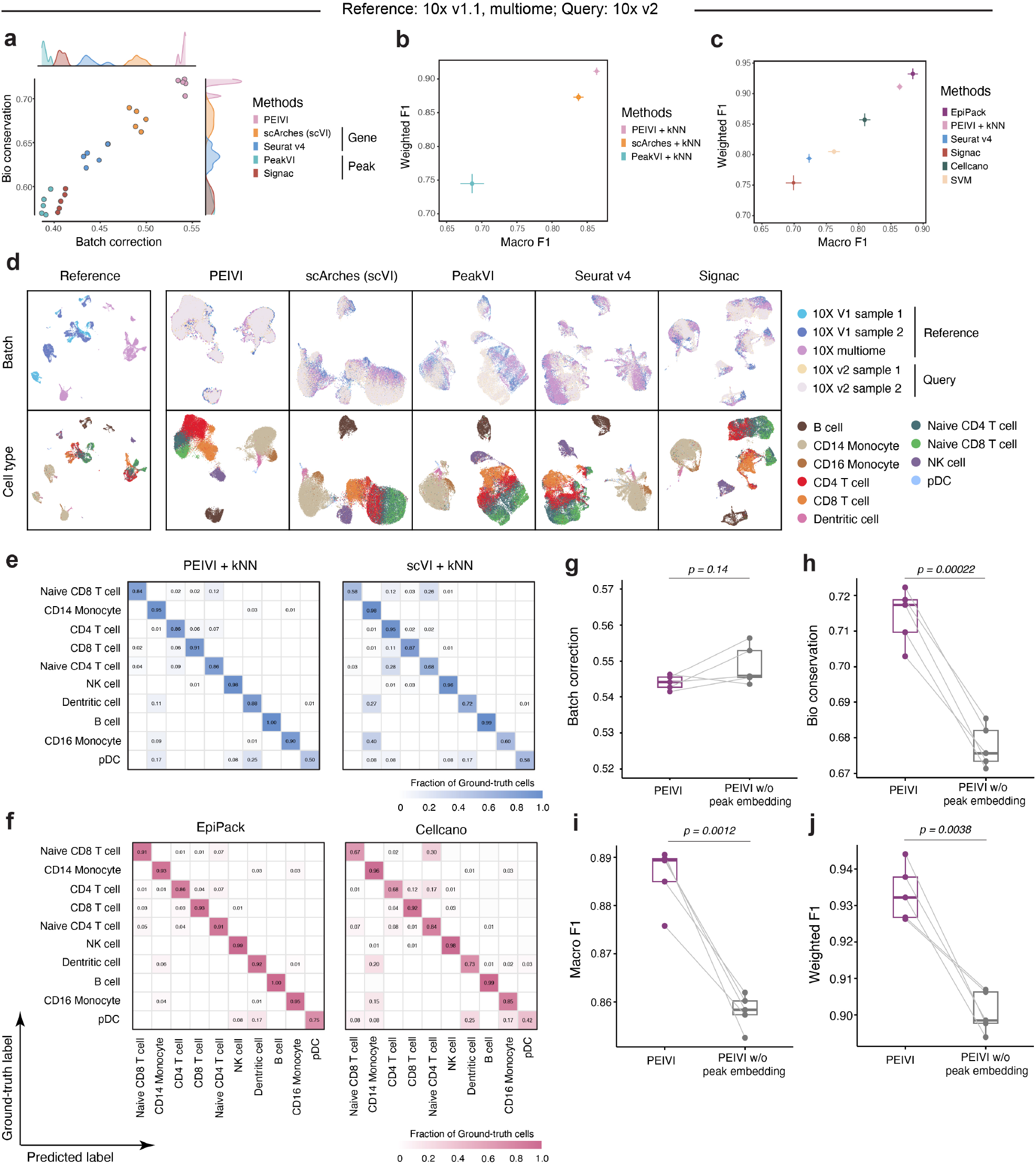
Accurate query mapping and label transfer with heterogeneous scATAC-seq features. **a**. Overall scores for the benchmarked models’ biological conservation and batch correction performance (n=5 for 5 repeating experiments). **b**. Cell label transfer performance using a kNN (k=5) on the joint embedding space across methods to reflect nearest neighbor structure preservation (n=5) in unsupervised reference mapping. **c**. Weighted F1 and Macro F1 scores of the benchmarked models in the supervised cell label transfer setting (n=5). **d**. UMAP visualization of reference mapping results. The top row shows batch labels; the bottom row shows cell type labels. **e**. Confusion matrices comparing PEIVI+kNN and scVI+kNN on cell type annotation. **f**. Confusion matrices comparing EpiPack and Cellcano classifiers. **g-j**, Ablation study comparing PEIVI with and without peak embedding (n=5). (g) comparable batch effect correction; (h) significantly higher biological conservation; and (i) improved macro and (j) weighted F1 scores.

High-quality mapping enables that query cell embeddings closely align with reference cell embeddings of the same type, meaning neighboring cells in the mapped space often share the same cell type annotation. Here we compared PEIVI with baseline models that also perform reference mapping, in terms of their performance in cell type annotation, by equipping each of them with a k-nearest neighbor (kNN) classifier to annotate query cell types in the joint embedding space after mapping. As evaluation metrics of the annotations, we used macro F1 and weighted F1 scores, where the former is more sensitive to minor populations, and the latter serves as a more balanced metric (Methods). We observed that PEIVI largely improved the resolution of the co-embedding space on scATAC-seq data by incorporating heterogeneous peak information. In the “Ref v1.1&multi, Query v2” group, PEIVI improved kNN label transfer accuracy by approximately 6% across both metrics compared to the second-best reference mapping method scVI+kNN (Fig.2b). By visualizing the confusion matrix of its annotation results (Fig.2e), we found that PEIVI achieves improved classification performance, especially when separating closely-related cell types. For example, for Naive CD8 and Naive CD4 cells, EpiPack increases the average classification accuracy from 0.58 to 0.84 and 0.68 to 0.86 respectively, compared to scVI, which is over 30% of improvement (Fig.2e). Similar performances can be found in the other two experimental settings (Supplementary Fig.3). All results indicate that using heterogeneous features including high-resolution peak information allows PEIVI to learn a better-aligned joint embedding space.

By replacing the kNN classifier with EpiPack’s metric learning-based classifier, EpiPack further improves classification performance. Compared to the plain kNN, the EpiPack Classifier excelled in cell type annotation tasks within the heterogeneous transfer space, particularly for underrepresented cell types, with the average macro F1 score improving from 0.86 to 0.89 (Fig.2c). Additionally, compared to other popular annotation tools for scATAC-seq, EpiPack consistently achieved higher weighted and macro F1 scores (Fig.2c). The confusion matrix clearly demonstrates EpiPack’s classification performance across all cell types (Fig.2f). Especially, for the classification of a minor population, pDC cells (with only 12 cells), EpiPack achieved an accuracy rate of 75%, while Cellcano, the second-best performer, achieved only 42%. When distinguishing CD16 monocytes from closely-related cell types like CD14 monocytes, EpiPack achieved accuracy rates of 0.95, while Cellcano reached only 0.85. Such improvements were also exhibited across the other two query groups (Supplementary Fig.4). The observations show that EpiPack achieves superior classification performance by integrating heterogeneous feature information with the metric learning classifier, including annotating rare cell types.

To determine whether the improvements arose from the incorporation of heterogeneous peak information rather than the PEIVI model design, we removed the peak constraint term (Eq. 14) and trained the PEIVI model solely using the gene score matrices as an ablation study. As expected, the results indicated that incorporating embedded peak information significantly enhanced the biological conservation performance of our model in the query mapping task (Fig.2h), while, compared to the plain gene score-based variant, it did not incur a significant loss in batch correction performance (Fig.2g). This finer-grained information transfer also improved the accuracy of cell type annotation. As shown in Fig.2i-j, PEIVI enhanced classifier performance in terms of both macro F1 and weighted F1 scores by incorporating peak information, further validating the advantage of the heterogeneous model in joint embedding space modeling.

### Leveraging heterogeneous features enhances cross-reference genome mapping and annotation performance

Due to the rapid iteration of reference genome versions, when utilizing previously published datasets as the reference atlas, scATAC-seq reference and query data may originate from different reference genomes. This discrepancy leads to greater feature space divergence. Such a challenge reduces the transferability of the foundational model for scATAC-seq data, highlighting the increasing importance of developing a reference genome-stable mapping approach. By leveraging heterogeneous features, PEIVI effectively preserves peak-level information, thereby providing a principled solution to overcome this issue. Thus we proceeded to perform this more challenging test of mapping and annotating datasets that are generated with different reference genomes. We used five PBMC datasets generated with Hg38 as the reference and one PBMC dataset generated with Hg19 as the query to benchmark PEIVI’s performance in cross reference genome mapping and label transfer tasks. Similarly, for methods that require aligned features, we performed benchmarking using gene scores and aligned peaks, where peak shifts caused by differences in reference genomes were corrected using Liftover^37^(Methods). The results showed that, compared to forcibly aligning feature spaces, leveraging a heterogeneous feature set led to superior performance in reference mapping tasks. Among these methods, PEIVI consistently outperformed other approaches that rely solely on homogeneous features (Extended Data Fig.2a). Compared to mapping and label transfer tasks within the same reference genome, differences in reference genomes deteriorated the performance of peak-alignment-based methods. In contrast, the heterogeneous transfer model effectively circumvented this limitation. Additionally, we visualized the joint embedding spaces of PEIVI and scArches, the second-best method, using UMAP. The results showed that incorporating heterogeneous features improved the alignment of cell populations with their nearest neighbors in the reference space (Extended Data Fig.2d).

The quantitative results of the cell label transfer task further demonstrated EpiPack’s superior classification performance compared to the gene score-based scArches (scVI) in both general transfer tasks and minor population transfer tasks (Extended Data Fig.2b,c), where a larger improvement is achieved in cross-reference genome annotation compared to the case of using the same reference genome. Specifically, it outperformed the second-best method, increasing the macro F1 score by approximately 13.9% (from an average of 0.66 to 0.75) and the weighted F1 score by approximately 15.8% (from an average of 0.76 to 0.90). Compared to the state-of-the-art method Cellcano, EpiPack achieved an improvement of over 20%. We also visualized EpiPack’s classification accuracy, as shown in Extended Data Fig.2e. The results demonstrated that, compared to scArches and Cellcano, EpiPack achieved significantly improved classification accuracy for low-abundance naive CD8 T cells, dendritic cells, and CD16 monocytes. These results, together with the benchmarking results under the same reference genome background, supported that the heterogeneous transfer model provides a more effective solution for scATAC-seq reference mapping and cell type annotation. Compared to homogeneous models, it not only significantly enhances model performance while improving scalability but also increases the efficiency of utilizing scATAC cell atlases published at different time points.

### EpiPack PEIVI builds effective and robust mapping space

An effective and mappable foundational model is a prerequisite for successful reference mapping and identifying cell types and states from the reference. While EpiPack has already demonstrated outstanding performance in mapping and cell annotation, we also recognized the need to evaluate its atlas construction capability. To demonstrate this, we benchmarked EpiPack PEIVI against various existing data integration methods using two gold-standard datasets with varying batch scales^8, 20, 38–41^. The mouse brain dataset includes two batches with approximately 10k cells, while the human PBMC dataset comprises five batches with around 40k cells (Methods). For fair comparison, all benchmarked methods were tested using their best-performing feature types^18^. Our findings, illustrated in Fig.3a and Supplementary Fig.5a, showed that PEIVI achieved performance comparable to, and in some cases surpassing, state-of-the-art methods. Supplementary Fig.6,7 show the UMAP visualizations of the aligned cell embeddings. Similarly, we also benchmarked PEIVI’s integration performance in the cross-reference genome scenario, where PEIVI demonstrated a significant enhancement in overall accuracy, as well as improvements in both biological conservation and batch correction performances when compared to other methods that use either remapped peak features or gene score features (Supplementary Fig.5b).

**Fig. 3.**
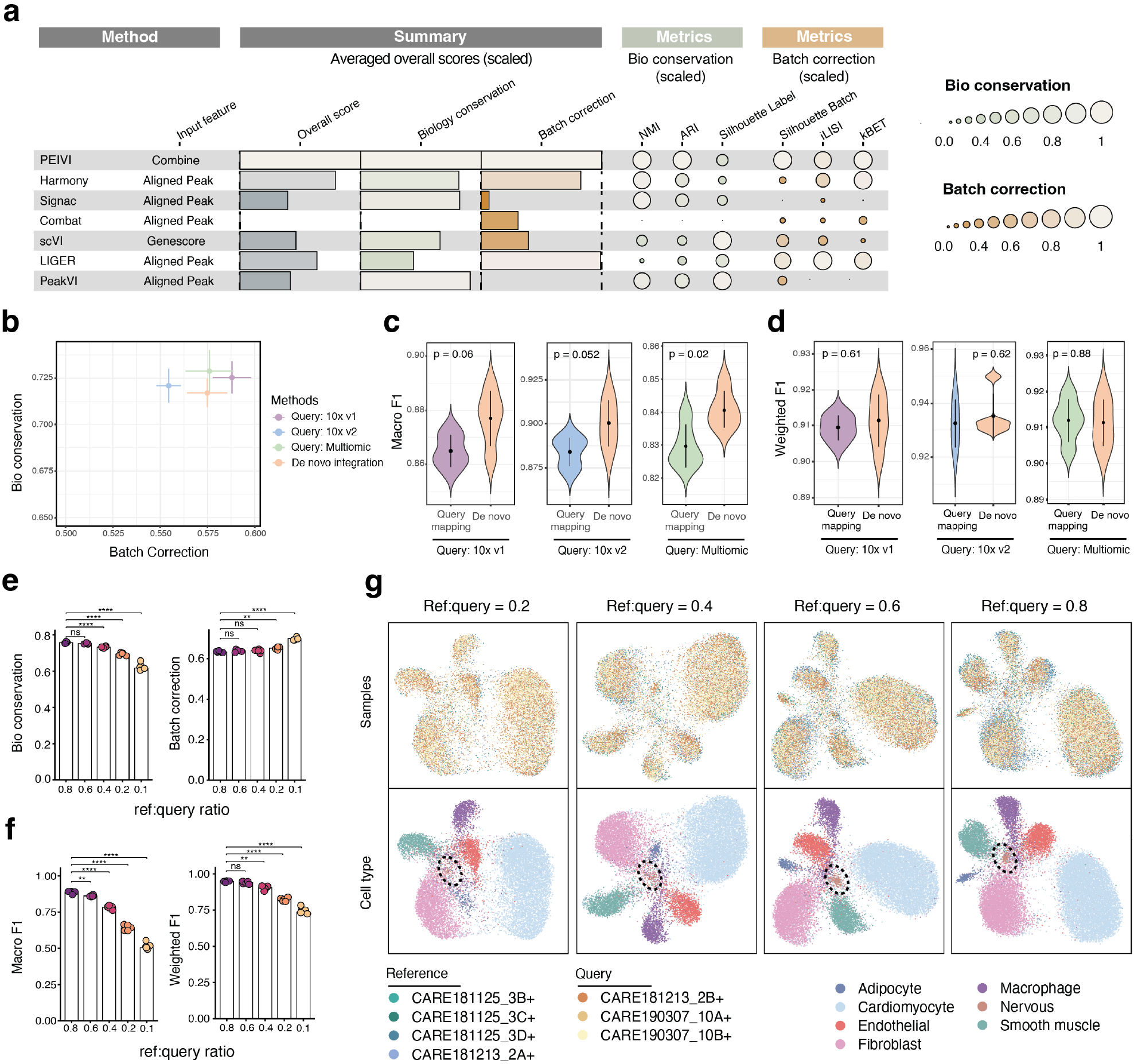
PEIVI constructs a robust and biologically meaningful reference mapping space. **a**. Benchmarking of PEIVI against six widely used data integration methods with different types of aligned input features on the human PBMC dataset. **b**. Comparison of PEIVI reference mapping versus de novo integration across different query groups. Comparable performances are shown in terms of biological conservation and batch correction. **c-d**. Violin plots comparing macro F1 (c) and weighted F1 (d) scores between PEIVI reference mapping and de novo integration across three query conditions. Results are based on five independent runs. Central dots indicate the mean; vertical lines represent ±1 standard deviation. P-values were calculated using two-sided unpaired t-tests and are shown above each comparison. **e-f**. Performance of PEIVI under varying reference-to-query ratios in the Cardiac Atlas dataset, based on five independent repeats. (e) Biological conservation and batch correction metrics decline significantly when the ratio drops below 0.6. (f) Macro and weighted F1 scores show decreased performance at lower reference sizes. Asterisks indicate statistical significance (two-sided t-tests, **** means p < 0.0001). **g**. UMAP visualizations of PEIVI mapping at different ref:query ratios. Distinct Nervous cell clusters (highlighted by black circles) emerge when the ref:query ratio exceeds 0.6.

We next investigated whether PEIVI reference mapping could approximate the performance of *de novo* integration. *de novo* integration is advantageous over reference mapping in terms of aligning difference batches because integration is performed from scratch each time there is a new batch, but is not scalable. In the benchmarking, we evaluated the performance using both mapping scores and classification metrics. For *de novo* integration, the classification metrics were calculated using datasets that corresponded to the respective query mapping data. Fig.3b shows that the overall performance of PEIVI reference mapping and *de novo* integration are similar. In particular, the biological conservation scores across all query groups closely align with the integration performance. Correspondingly, the classifier’s cell type annotation performance in the reference mapping space showed no significant difference from its performance in the *de novo* joint embedding space (Fig.3c,d). This further validates that the mapping space learned by PEIVI effectively preserves its biological characteristics.

As a deep transfer learning model, PEIVI’s query mapping performance is typically influenced by the scale of the reference data, with smaller datasets potentially leading to underfitting. To assess the model’s sensitivity to data scale, we examined reference mapping and cell label transfer performance under different reference-to-query ratios. We leveraged the Cardiac Atlas datasets^2^for benchmarking, selecting four samples as reference data and three samples as query data. By progressively downsampling the reference data, we adjusted the ref:query ratio and constructed PEIVI models under 80%, 60%, 40%, 20%, and 10% reference-to-query conditions. We then quantified mapping and annotation performance across these reference models. In the test, we observed that as the ref-query ratio dropped below 0.6, the model’s mapping effectiveness and cell type transfer capability significantly declined. (Fig.3e,f). Notably, from UMAP visualizations, we observed distinct clusters of neuronal cells only when the reference-to-query ratio exceeded the 0.6 threshold. In contrast, when the ratio fell below this threshold, these cells became indistinguishable from the background within the mapping space (Fig.3g). We repeated the test on PBMC datasets, which exhibited similar performance (Supplementary Fig.8). Meanwhile, we also noticed that the reference mapping and cell type annotation performance of EpiPack remains robust across various hyperparameter settings (Extended Data Fig.3,4).

Given the inherent imbalance in single-cell data^42^, accurately labeling the extremely rare cell populations (less than 1%) can be challenging^42^. We then investigated EpiPack’s cluster separability on rare cell populations at different proportions (Methods). We designed two test cases: (1) a scenario where the rare cell type close to one of the major cell types, achieved by downsampling CD16+ monocytes (which is close to CD14+ monocytes), and (2) a scenario where the rare cell type is distinct from all other cell types, tested by downsampling B cells (Supplementary Fig.9). The target cell types were downsampled to constitute 0.3%–1% of the whole dataset. The results indicate that in the CD16+ monocytes group, which shares similarities with major cell types, the rare cells form a clear cluster once their proportion reaches 0.5% of the whole dataset. In contrast, for B cells, which exhibit higher independence from other cell types, PEIVI maintains a stable joint embedding space when cell proportion varies.

### PEIVI preserves discrete OOR cell clusters and continuous cancer-associated immune cell dynamics within the mapping space

Since cell types and states in disease samples can shift due to perturbations, the query dataset may not have exactly the same cell types as in the reference dataset. A mapping algorithm should not only map identical cell types from both the reference and query datasets in the joint embedding space, but also preserve novel OOR (out-of-reference) cell types and states. Therefore, we proceeded to evaluate PEIVI’s performance in mapping such perturbed (OOR) cell types or states that are not present in the reference.

We first demonstrated PEIVI’s ability to preserve discrete OOR cell types (global OOR). To achieve this, we take the human PBMC dataset and split data from one sequencing platform as a pseudo-disease query dataset while using the remaining data as a reference control. Specifically, we removed one cell type (e.g., B Cells) from the reference dataset that was present in the query, ensuring that this cell type existed only in the pseudo-disease dataset. Based on this setup, we trained PEIVI where B cells were absent from the reference atlas. After fine tuning, in the integrated joint latent embedding space, we observed an independent B cell cluster, indicating that PEIVI effectively maintained the separability of OOR cell types. Repeated experiments across all sequencing platforms produced consistent results (Fig.4a-c). Next, we iteratively removed each cell type to construct pseudo-disease datasets with varying OOR population sizes and quantified the distinctiveness of OOR clusters using cell-type-specific ASW scores. The results demonstrated that our model consistently preserved and effectively separated OOR cell populations, regardless of their population size (Fig.4d) .

**Fig. 4.**
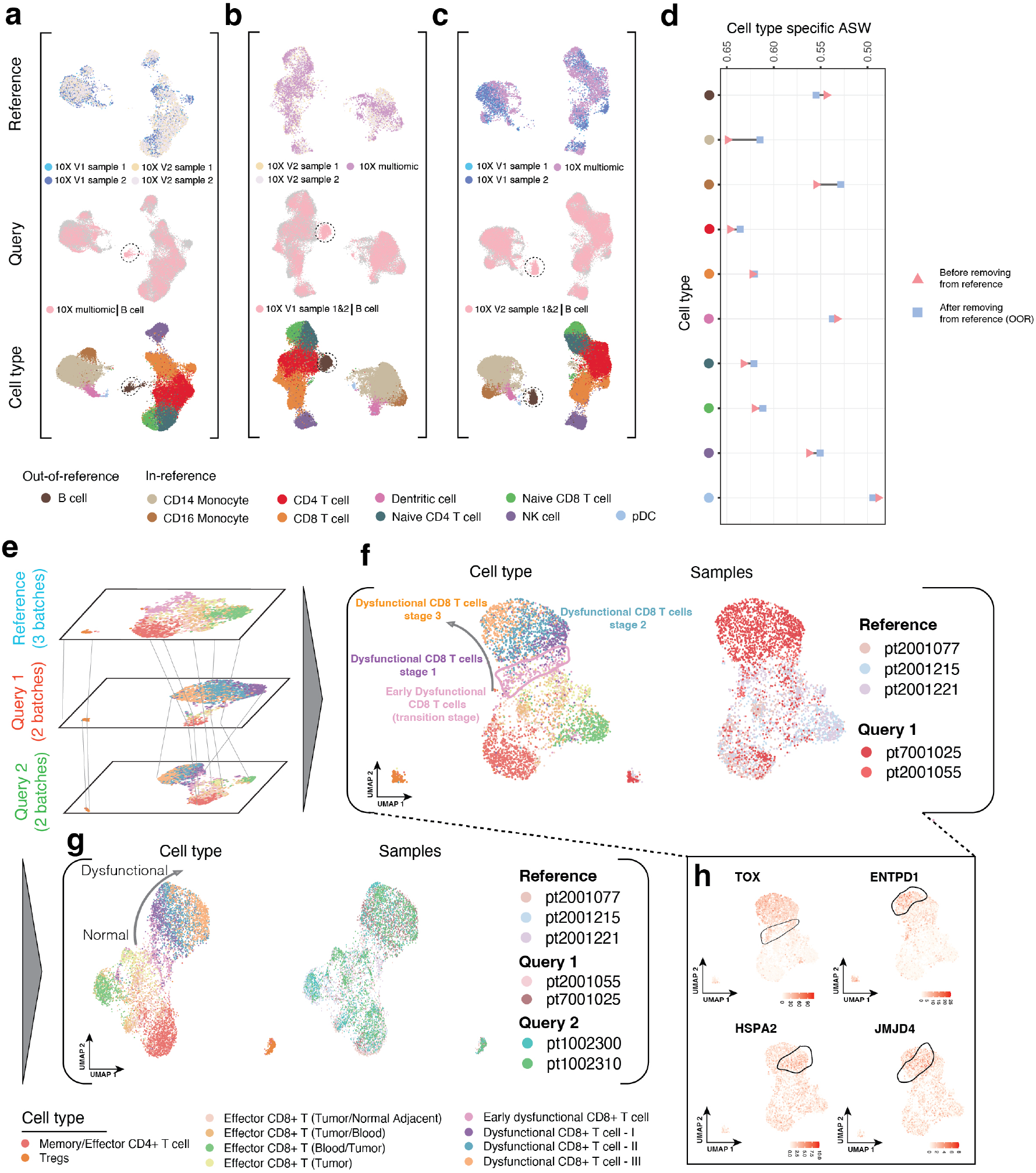
PEIVI preserves discrete cell localization and continuous immune cell state transitions in scATAC-seq query mapping. **a-c** UMAP visualizations of reference mapping in synthetic pseudo-disease settings, where one cell type (B cells, highlighted by black circle) is removed from the reference but retained in the query to simulate global out-of-reference (OOR) cell types. **d**. Dumbbell chart showing cell type-specific average silhouette width (ASW) scores before (triangles) and after (circles) removing the corresponding cell type from the reference. Higher ASW indicates better cluster separability. **e**. Schematic of the experimental setup using scATAC-seq data from ccRCC, where the reference and two query sets contain different subsets of CD8 T cell states. **f**. Query 1 mapping space: PEIVI preserves the full trajectory of dysfunctional CD8 T cells, including early, intermediate, and late stages, forming a continuous manifold with early dysfunctional cells in the reference. **g**. In Query 2, built upon a reference atlas that includes Query 1, cells are precisely mapped to their expected positions along the dysfunction transition gradient. **h**. UMAP projections of gene activity scores for stage-specific marker genes (*TOX, ENTPD1, HSPA2*, and *JMJD4*) validate the accurate positioning of query cells along the dynamic trajectory of CD8 T cell dysfunction.

Next, we evaluated PEIVI’s ability to position cells along chromatin dynamic gradients, which typically occur during continuous cell activation or dysfunction. In this test, we utilized scATAC-seq data from tumor tissues of seven early-stage clear cell renal cell carcinoma (ccRCC) samples^43^. We first constructed a reference atlas using three samples containing normal CD4 and CD8 T cells as well as early dysfunctional CD8 T cells. Next, we designated two distinct sets of samples with varying cell populations as query datasets: the first set primarily contained remaining gradients of dysfunctional CD8 T cells as the OOR cell states (Query 1), used to assess PEIVI’s mapping capability in positioning cells along a continuous cellular manifold; the second set encompassed all cell types and states (Query 2), intended to evaluate PEIVI’s performance in mapping existing cellular gradients (Fig.4e). All ground-truth cell labels were provided along with the original dataset.

In the mapping space of Query 1, we observed that the developmental trajectory of dysfunctional CD8 T cells was well preserved, forming a continuous manifold with Early dysfunctional CD8 T cells, consistent with previously published findings^43^(Fig.4f). The transition from early to late-stage Dysfunctional CD8 T cells was marked by the gradual expression changes of key genes identified by Kourtis et al., where *ENTPD1* was predominantly expressed in late Dysfunctional CD8 T cells, *JMJD4* expression increased during the mid-to-late dysfunction transition, and *HSPA2* exhibited relatively high expression levels in primary Dysfunctional CD8 T cells. The gene accessible score patterns of Early dysfunctional CD8 T cells resembled those of stage 1-3 cells, albeit with lower intensity (Fig.4h). These gradual expression patterns highlight the accurate positioning of OOR cell states along the transition gradient, reinforcing PEIVI’s capability to capture cellular dynamics within the mapping space.

Subsequently, in the mapping space of Query 2, which was mapped on the base model integrating the original reference with Query 1, query cells were accurately mapped to their corresponding cell state positions, with the dysfunction transition gradient also precisely aligned (Fig.4g). Using the EpiPack classifier, we inferred cell states and validated the accuracy of cell type assignments based on ground-truth labels, achieving a weighted F1 score of 0.70 and a macro F1 score of 0.71. The visualization of predicted labels against ground-truth annotations demonstrated precise positioning along the continuous transition gradient (Supplementary Fig 10), confirming the robustness of PEIVI’s mapping performance on the continuous cell dynamic pattern.

### Fast and precise OOR detection by EpiPack in pseudo-perturbation settings

The ultimate goal of EpiPack is not only to transfer cell type labels from the mapping space, but also to identify out-of-reference (OOR) cell types and states and return associated uncertainty quantification. Such OOR detection can provide valuable insights into perturbed cell state compositions under disease conditions. After showing PEIVI’s capability to preserve both discrete and continuous perturbed cell types and states, we now investigate whether EpiPack’s OOR detector can effectively identify OOR populations from the query data. To enable benchmarking under conditions with ground-truth labels, we continued to use the human PBMC dataset to simulate the test data. Two samples were selected as the pseudo-perturbation group, while the remaining three were used as the reference control.

We first simulated discrete OOR populations (global OOR types) by selecting a distinct cell type and removing all cells with that label from the reference group (Fig.5a). The resulting pseudo-perturbation dataset was mapped onto the pre-trained control reference latent space. Since the EpiPack global OOR detector is inherently coupled with its classifier, we benchmarked its performance against two widely used alternatives for novel cell type detection within classification space, namely a k-nearest neighbor (kNN) detector and a support vector machine (SVM) detector^12, 44^, both of which output probabilistic scores that can be thresholded to identify OOR cells. To ensure comparability, all methods were evaluated using the same integrated latent space from PEIVI as input. When B cell (which is a cell type well separated from other cell types) and CD8 T cell (which is close to other cell types) populations were removed from the reference dataset, respectively, EpiPack consistently achieved higher sensitivity and more effective FDR control across parameter settings (Fig.5b, Supplementary Fig.11). We further compared OOR detector performance on PEIVI embeddings with that on scArches (scVI) embeddings based on homogeneous gene score features, and found that PEIVI-based detectors maintained higher true positive rates (TPR) while substantially reducing false discovery rates (FDR), providing more robust overall performance. The improvement was most pronounced for the CD8 T cell population (∼ 23.6%), but was also evident for the more distinct B cell population (∼ 6%) (Fig.5c). We attribute this to the fact that global OOR detection relies on clear separation between clusters, which is achieved by PEIVI through incorporating peak-level priors, thereby enhancing sensitivity while reducing misclassification of in-reference cells as OOR.

**Fig. 5.**
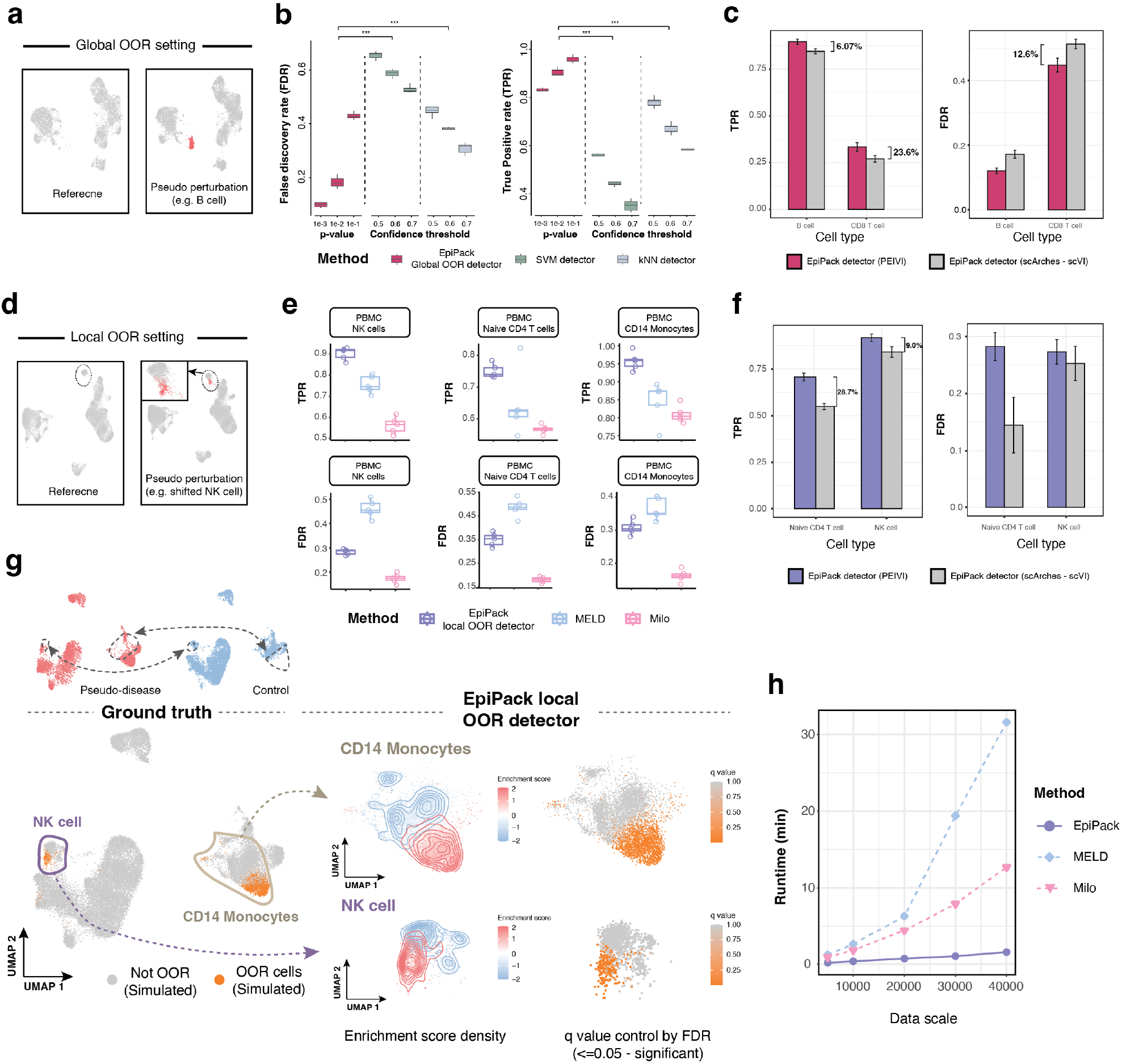
Fast and precise detection of out-of-reference (OOR) cell types and states by EpiPack. **a**. Global OOR simulation by removing a distinct population (e.g. B cells) from the reference. **b**. Boxplot shows that EpiPack global OOR detector achieves higher sensitivity and stronger FDR control than kNN and SVM classifiers on B cells (n=5 for 5 repeating experiments, two-sided t-tests, **** means p < 0.001). **c**. Comparison of PEIVI-versus scVI-based embeddings reveals that PEIVI substantially improves detection accuracy (n=5). **d**. Local OOR simulation generated by shifting subsets of the selected cell population (e.g. NK cell) in latent space. **e**. Boxplots show that across different PBMC cell type backgrounds, the EpiPack detector achieves higher sensitivity and maintains FDR control compared with benchmarked methods. **f**. Embedding choice impacts local detection, with PEIVI enhancing sensitivity but elevating FDR relative to scVI. **g**. Visualization of mixed perturbations demonstrates that EpiPack accurately localizes OOR state shifts, with enrichment scores and q-values aligning with ground-truth regions.**h**. Line plot of running time for EpiPack, Milo, and MELD across different atlas scales.

We next evaluated the performance of the EpiPack local OOR detector in discovering perturbed cell state shifts. To simulate local OOR states in pseudo-disease datasets, we selected one annotated cell type and removed a fraction of those cells from both the reference control and pseudo-perturbation datasets along the principal component axis, thereby generating controlled OOR state shifts with ground-truth labels (Fig.5d, Methods). Following the rationale of Dann et al.^45^, directly detecting OOR states from an atlas reference alone can introduce unstable biases. Therefore, in this scenario, we constructed a harmonized latent embedding by jointly integrating all reference and perturbed datasets, rather than relying on reference mapping, and subsequently benchmarked detection methods using this embedding. To benchmark EpiPack’s local OOR detector, we compared it against Milo^24^and MELD^26^, two clustering-free methods that operate directly in latent space for OOR state detection.

EpiPack local OOR detector demonstrated high sensitivity in detecting simulated OOR state regions while maintaining stable FDR control across benchmarking scenarios (Fig.5e). Across different cell type backgrounds, our method consistently achieved the highest sensitivity. In contrast, MELD exhibited poor FDR control due to the absence of statistical testing and the oversmoothing introduced by its fixed-kernel design, whereas Milo provided more balanced performance, with better FDR control but substantially lower sensitivity compared to EpiPack. We next compared the embedding from PEIVI and scVI in terms of local OOR state detection. Interestingly, in the local OOR setting (Fig.5f), while PEIVI again achieved higher TPR, it also exhibited an elevated FDR, indicating a more sensitive but less conservative behavior relative to scVI. Together with its performance in global OOR detection (Fig.5c), we consider that this is because local OOR detection depends critically on the local manifold geometry and continuity of cellular states, and PEIVI enables subtle shifts to be captured by preserving fine-grained heterogeneity in the embedding, thereby improving sensitivity (TPR), but at the cost of misclassifying some residual noise or batch effects as OOR, which elevates FDR.

In addition, the EpiPack detector demonstrated superior computational efficiency (Fig.5h). To ensure a fair comparison, all methods were benchmarked on a CPU platform (AMD EPYC 32-Core Processor). Compared with MELD and Milo, EpiPack substantially reduced runtime by applying a fixed pre-trained kernel for diffusion across the entire graph. Within datasets ranging from 5,000 to 40,000 cells, EpiPack consistently achieved the fastest performance, lowering runtime costs to approximately 10% of the baseline (Milo). This efficiency markedly enhances its applicability to atlas-scale datasets.

We next visualized the performance of the EpiPack local OOR detector under a mixed OOR state shift scenario, in which both NK cells and CD14 monocytes were simultaneously perturbed (Fig.5g). This setting was designed to assess whether the detector could remain robust in the presence of multiple concurrent shifts. The results indicate that under this complex perturbation, EpiPack outputs q-values that align with the ground-truth OOR states. In summary, these experiments demonstrate the sensitivity of the EpiPack OOR detector in both global and local OOR detection contexts.

### EpiPack discovered disease-related immune populations in COVID-19 dataset

To demonstrate the utility of EpiPack in identifying potential perturbed cell populations through reference mapping under disease contexts, we collected published scATAC-seq datasets of PBMCs from nine healthy donors and patients^46^. As the reference atlas, we harmonized 75,193 healthy PBMC cells to pre-train the base model and annotated 10 cell types using the same strategy as applied in the PBMC benchmark dataset (Fig.6a, Supplementary Fig.12). Following the “Atlas to control reference” OOR detection experiment strategy recommended by Dann et al.^45^, we jointly mapped healthy control PBMCs (n = 17,258) and COVID-19 PBMCs (n = 13,205) to the atlas reference and performed label transfer using the EpiPack classifier (Fig.6b, Methods). To ensure the reliability of the annotations, we additionally visualized key marker genes(Supplementary Fig.13), which confirmed expected gene score distributions.

**Fig. 6.**
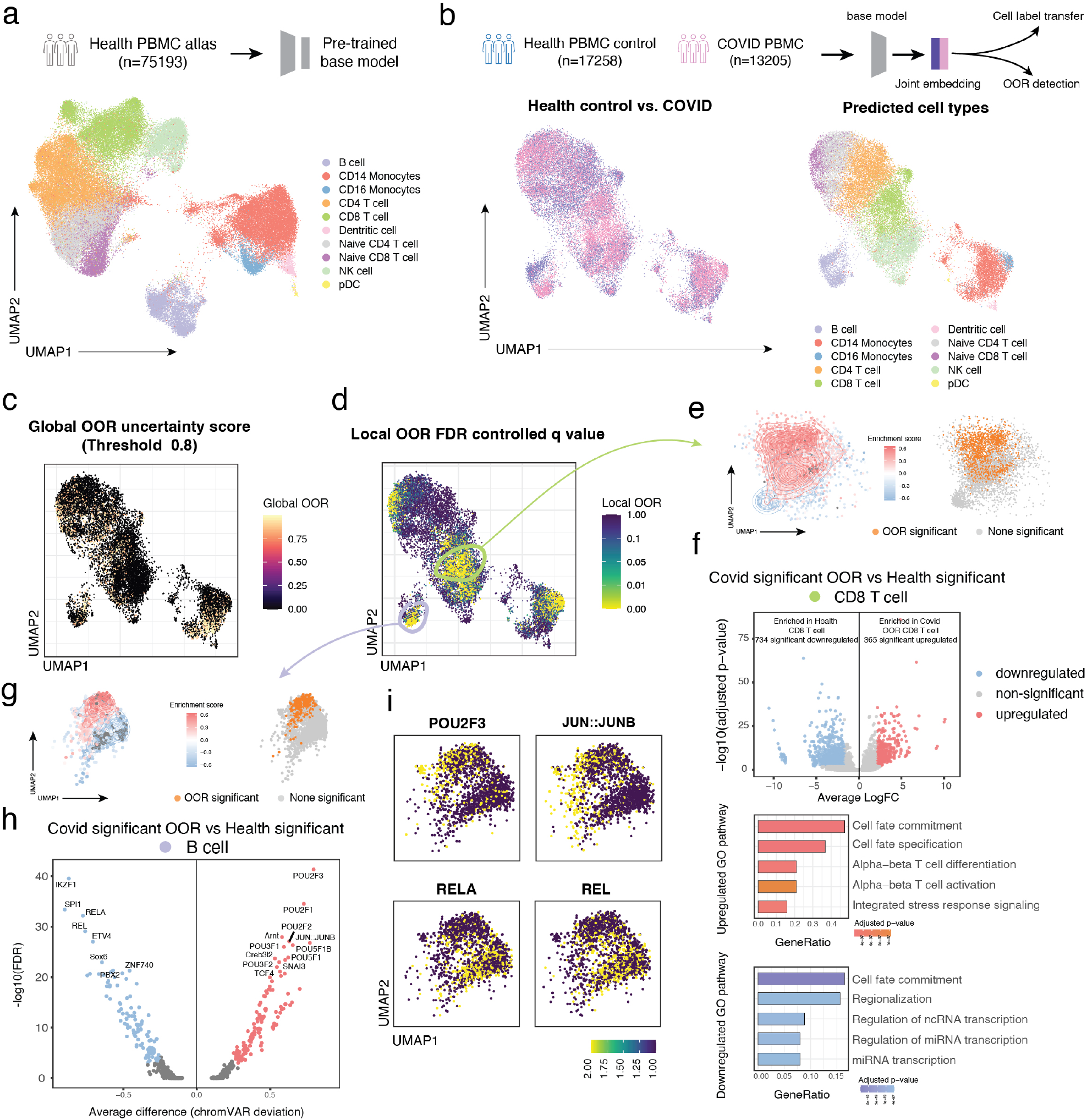
EpiPack identifies disease-associated CD8 T cell and B cell populations in COVID-19 PBMCs. **a**. UMAP visualization of the reference atlas of 75,193 healthy PBMCs used for pre-training the base model with 10 annotated immune cell types. **b**. Joint mapping of healthy control (n = 17,258) and COVID-19 (n = 13,205) PBMCs to the atlas with label transfer. **c**. UMAP visualization of the global OOR detection. All cells with an uncertainty score < 0.8 were set to 0 to facilitate visualization of significant OOR clusters. **d**. UMAP visualization of the global OOR detection. **e**. UMAP visualization of enrichment scores (left) and significant OOR cells (right) within CD8 T cells. **f**. Top: Volcano plot of differential peak analysis between healthy and COVID-associated CD8 T cells. Peaks with logFC > 2 or < −2 and −log10(adjusted p-value) > 3 were defined as significant. Down: Bar plots displaying the most significantly enriched GO biological processes associated with differential peaks in each cluster. **g**. UMAP visualization of enrichment scores and significant OOR cells within B cells.**h**. Volcano plots showing differential TF motif accessibility, based on mean chromVAR bias-corrected deviation scores, between healthy and COVID-associated OOR B cell states. **i**. UMAP visualization of the selected TF regulator activity

The UMAP visualization in Fig.6b presented a clear distributional shift between the COVID-19 and healthy control groups. We therefore applied the EpiPack global and local OOR detectors to identify cell types and states in the COVID-19 group that may represent perturbations relative to healthy controls. We first applied the global OOR detector directly to the healthy control and COVID-19 datasets to assess whether any discrete cell clusters with significant shifts were present in the COVID-19 samples. The results showed that the global detector did not identify distinct OOR clusters, suggesting that no clearly segregated novel cell types emerged in this cohort (Fig.6c). However, the global detector also highlights subpopulations with low-confidence during label transfer within the CD8 T cell, B cell, and naive CD8 T cell regions, implying potential cluster shifts in these compartments. We next applied the local OOR detector to the COVID-19 group, where FDR-controlled q-values revealed patterns of COVID-associated cellular shifts. Regions with q-values < 0.05 were primarily localized to CD8 T cells, naive CD8 T cells, B cells, and CD16 monocytes (Fig.6d), consistent with the regions highlighted by the global OOR detector. Given their central role in antiviral immunity^47, 48^, we next focus on CD8 T cells and B cells, where perturbation signals were most prominent.

We first visualized in detail the enrichment scores and significant OOR cells within the CD8 T cell population. As expected, a substantial fraction of these cells from the health and COVID-19 conditions form partially overlapping polarized manifolds, with regions of elevated enrichment scores coinciding with those predicted as OOR cells (Fig.6e). To determine whether these detected OOR cells represent disease-associated phenotypes and to define the transcription factor (TF) signatures underlying the COVID-associated CD8 T cell response, we performed differential peak analysis between OOR cells and highly enriched healthy counterparts (Methods). In CD8 T cells, we identified 19,205 differentially accessible peaks, of which 365 were significantly upregulated and 734 were significantly downregulated (Fig.6f). These significant peaks were further searched against a panel of cell type–differential peaks to identify overrepresented DNA motifs, followed by GO analysis of the enriched motifs (Methods). The results revealed a major upregulation of functional programs associated with cell fate commitment as well as αβ T cell differentiation and activation within the OOR population, indicating a shift toward rapid effector responses under inflammatory conditions. Collectively, these findings suggest that the CD8 T cell OOR cluster represents a disease-induced activated subpopulation, consistent with findings on CD8 T cell activation reported in previous studies^48–50^.

In B cells, we similarly observed that OOR cells coincided with regions of high enrichment scores (Fig.6g). Differential accessibility analysis identified 18,472 peaks, of which 210 were significantly upregulated and 141 were significantly downregulated (Supplementary Fig.14a). GO analysis revealed upregulation of pathways related to miRNA regulation, alongside downregulation of gland development–associated programs (Supplementary Fig.14b). To further characterize regulatory heterogeneity, we assessed TF deviation scores and variance between B cell subpopulations from enriched healthy cells and detected COVID-associated OOR cells. Notably, NF-κB subunits involved in germinal center B cell maintenance (e.g. REL and RELA)^51, 52^were enriched in the healthy B cell subpopulation but relatively silent in the OOR B cells cluster(Fig.6h,i), consistent with the GO pathway analysis and reflecting impaired germinal center function in COVID-19^53^. In contrast, transcription factors involved in B cell receptor signaling and indicative of B cell differentiation and activation, including JUNB and POU2F2/3^54^, were enriched in the detected OOR B cell subpopulation (Fig. 6h,i). These findings suggest that the OOR B cell cluster represents a COVID-induced activated B cell population.

Overall, EpiPack, through reference mapping and a multi-stage OOR detection framework, effectively captures biologically relevant variation signals and uncovers TF motif alterations of functional significance.

## Discussion

In this study, we present EpiPack, a comprehensive deep learning framework for scATAC-seq reference atlas construction, query mapping, cell label transfer, and out-of-reference (OOR) detection. Central to EpiPack is the Peak Embedding Informed Variational Inference (PEIVI) model, which leverages the paradigm of heterogeneous transfer learning to support reference model construction and unsupervised query mapping across varying sequencing protocols and reference genome backgrounds. Building on this foundation, EpiPack introduces a mathematically grounded global–local OOR detection framework, which formally distinguishes discrete and continuous forms of OOR cell types or states, and is equipped with classifiers tailored to their respective metric spaces. Comprehensive benchmarking demonstrates that PEIVI’s unique design outperforms existing methods based on homogeneous features in both data integration and transfer tasks for scATAC-seq, while also mitigating challenges introduced by reference genome version variation in real-world analyses. Additionally, the EpiPack OOR detector provides a more effective and interpretable solution than current models for detecting potential perturbed cell types. In practice, EpiPack supports a full analysis workflow—from pretraining on labeled references to model deployment, from annotating new data to quantifying OOR uncertainty, and from identifying discrete OOR cell populations to detecting continuous perturbations—making it broadly applicable across a range of foundational scATAC-seq analysis scenarios.

Reference mapping in scATAC-seq shares considerable conceptual similarity with that in scRNA-seq, but the unique structure of the scATAC-seq feature space presents greater challenges for feature alignment^55–57^. In fact, the choice of feature space has long been a fundamental issue in query-based data integration. While transforming peak features into gene-centric representations may appear to be a straightforward solution^12, 13^, such conversions inevitably incur information loss, which can be detrimental. Our experiments confirmed that alignment-based features led to a 10–20% performance drop compared to the heterogeneous peak embedding space introduced in our framework. During training, we adopted a gene score–based bridging strategy, which effectively reduced computational complexity by significantly lowering the dimensionality of input data compared to peak or fragment matrices. The lost information was then approximated and captured by incorporating peak information in the latent embedding space via learned constraints. Algorithmically, this approach can also be interpreted as a multi-view learning scheme^58^per sample—where the peak embedding view is used to reconstruct the gene score view. Consequently, the method also inherits limitations of this structure: its representational efficiency remains influenced by the information richness of gene score data, although this limitation is mitigated by the incorporation of peak embedding information.

Based on that, we envision potential directions for further improving our model. In this study, PEIVI primarily employs an autoencoder-based framework that operates on feature matrices, without directly leveraging sequence-level information from scATAC-seq fragments. Although the heterogeneous transfer model partially alleviates discrepancies arising from feature heterogeneity, its performance remains constrained by the quality of gene scores and thus cannot fully overcome the limitations of heterogeneous features. With the rapid advancement of large language models^59, 60^, particularly their enhanced capacity for long-sequence modeling, it becomes feasible to pre-train on informative fragment sequences and subsequently fine-tune on query data for tasks such as cell type annotation or zero-shot learning. Such an approach could mitigate the information loss inherent to feature transformation, offering a pathway toward more expressive and sequence-aware reference mapping in scATAC-seq analysis.

We also note that for the critical task of OOR detection in query mapping, our proposed global–local OOR detection framework provides a clear and biologically grounded delineation between two distinct classes of perturbation: those involving substantial biological divergence and those reflecting gradual state transitions. In particular, compared with previous approaches, our local OOR detector achieves markedly higher sensitivity by employing a trainable kernel and a bidirectional residual diffusion process that mimics the propagation of attention signals across subpopulations with distinct topological features. Moreover, by decoupling the attention kernel from the signal propagation process, the method substantially accelerates computation and reduces time complexity. More generally, because our OOR detector operates on a joint embedding rather than relying on an end-to-end architecture, it can also be applied to integrated scRNA-seq spaces, thereby extending its applicability across modalities.

In the COVID-19 case study, we demonstrated EpiPack’s application to disease data. EpiPack successfully identified COVID-associated OOR populations and uncovered perturbed motif patterns through region enrichment analysis. Such analyses can likewise be extended to other disease contexts to reveal biologically meaningful regulation of cis-regulatory elements.

We view EpiPack as a modular analysis platform for scATAC-seq data that makes three key contributions: (1) it provides valuable insights into the challenge of heterogeneous feature spaces in scATAC-seq data integration and mapping, offering a solution that preserves peak-level information; (2) its modular design and reusable reference model interface enable the effective utilization of large-scale pre-trained models for scATAC-seq; and (3) it introduces the first OOR detection framework for scATAC-seq, whose high sensitivity and computational efficiency extend the applicability of scATAC-seq to atlas-scale perturbation and disease analyses, thereby enriching the methodological toolkit for this modality. Looking forward, we anticipate that an increasing number of atlas-scale datasets will be pre-trained with EpiPack and shared through open-access repositories for community use. With the availability of such reference models, EpiPack is expected to accelerate the adoption of reference mapping in scATAC-seq, facilitating more efficient and scalable analyses of emerging datasets.

## Methods

### EpiPack toolkit architecture

EpiPack architecture can be briefly divided into two parts: (1) reference construction and mapping and (2) out-of-reference cell detection. The key idea of the EpiPack model is that the latent embedding generated by the gene score matrix can be learned to become a higher-resolution embedding result guided by the latent space of its corresponding peak, thereby overcoming the problem of scATAC-seq data feature distinction and obtaining more flexible reference model, which can be used to project query data on the reference embedding easily and transfer cell labels. On the basis of high-quality co-embedding, OOR cell types and states can be detected. In the following sections, we first provide a detailed explanation of the peak embedding-informed generative process, which represents one of our primary contributions. Then, we describe how cell discovery operates within the mapping embedding space, incorporating both local and global OOR algorithms.

### The peak embedding-informed variational inference (PEIVI) model

Assume that there are *J* scATAC-seq datasets from different batches or sources *b*_1_, *b*_2_, · · ·, *b*_*J*_ to be integrated, each with a distinct feature (peak) set *P*_*n*_, *n* = 1, 2, · · ·, *J*. To integrate and map scATAC-seq datasets characterized by distinct feature sets, we reason that a common latent space is required to locate different batches. As described previously, we utilize the gene score matrix as an effective bridge to link the peak embedding spaces of different batches. Thus, before integrating, we calculate a gene activity score matrix for each scATAC-seq batch *b*_*j*_, which is defined as *G*_*j*_ := *gene*_*score*(*b*_*j*_). The original gene activity score function *gene*_*score*(·) in EpiPack is based on the GeneActivity function in Signac^36^. Meanwhile, the gene score matrix can also be generated using snapATAC2’s make_gene_matrix function. Thus for each dataset *b*_*j*_, we have *{N*_*j*_, *P*_*j*_, *G*_*j*_*}* to denote *N*_*j*_ cells from the *j*th dataset with two matrices: binary peak count matrix 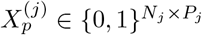 and gene score matrix 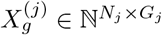.

To better preserve peak information and facilitate its transfer, we reason that a lower-dimensional embedding could be a suitable choice. Therefore, given the heterogeneity of feature sets, each batch is equipped with an independent dimension reduction unit that generates a batch-specific latent space, for which we use an autoencoder structure similar to BinaryVAE^61^(Fig. 1b step 1). We first model the binary peak count matrix from each scATAC-seq dataset as a low-dimensional latent variable vector, referred to as cell embedding 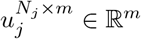, by a binary autoencoder, where *m* is the number of latent dimensions. We have

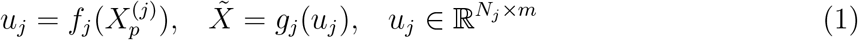

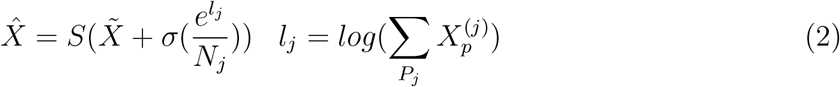

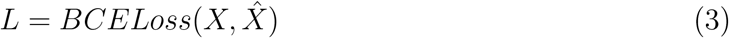

in which *f*(·) and *g*(·) are encoder and decoder functions respectively. *S*(·) is the sigmoid function in the output layer of *g*(·). *σ*(·) is the logit function that normalizes the library size^61^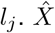 is the reconstructed gene expression matrix. The model is fitted by the binary cross-entropy

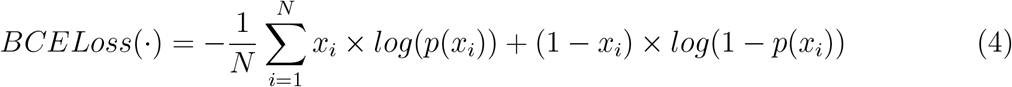

Considering that the peak autoencoders are trained independently using heterogeneous scATAC-seq datasets, it becomes crucial to establish a proper linkage between the cell embeddings learned by these autoencoders. Leveraging the inherent connection between the gene score matrix and peak count matrix in the same dataset and the common space provided by the unified gene score matrix, we propose Peak Embedding-Informed Variational Inference (PEIVI) model that uses a unified gene score matrix to link batch-specific latent spaces. Hereby, EpiPack *PEIVI* takes the gene score matrix 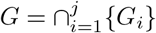 and peak cell embedding *U* = *{u*_1_, *u*_2_, · · ·, *u*_*j*_*}* as two inputs to link the autoencoders (Fig. 1b step 2). The gene score matrix is assembled using a shared gene set. Because the gene score matrix *G* also represents the cell identity to a certain extent (though not as much as the peaks), PEIVI should also be trained on *G* to generate the latent factor matrix *Z* = *{z*_1_, *z*_2_, · · ·, *z*_*j*_*}*, in which

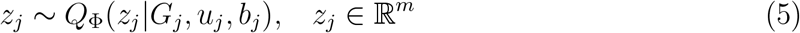

Then the goal of the model is to modify the latent embedding *Z* in the peak embedding-informed generative process. The generative process for observed data *G* involves maximizing the log-likelihood function log ℙ_Θ_(*G*|*U, B*) = Σ*j* log ℙ_Θ_(*G*_*j*_|*b*_*j*_, *u*_*j*_), where *θ* encompasses all decoder parameters. Since directly maximizing the log-likelihood requires integration over all possible *Z* values, which is computationally intractable, we employ a Variational Bayes approach.

For the sake of clarity, we will first elaborate on the scenario of training a standard VAE on G. In a standard VAE, the model will utilize the provided *Z* ∼ *Q*_Φ_(*Z*|*G, B*) as an approximate posterior distribution. Commonly, this distribution is modeled as a multivariate Gaussian. The optimization objective, referred to as the evidence lower bound (ELBO), is defined as:

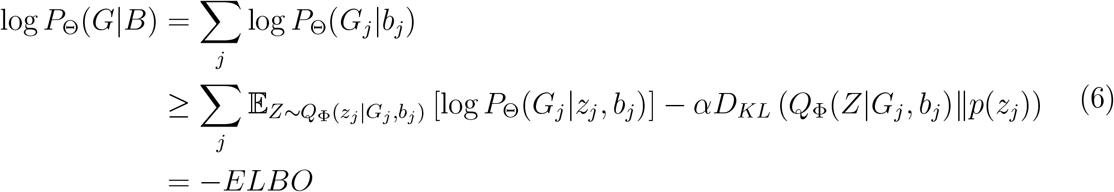

in which *Q*_Θ_(·) and *P*_Φ_(·) denote the encoder with parameter Φ and the decoder with parameter Θ respectively. *p*(*z*) represents the prior distribution of *z* which follows a standard multivariate normal distribution *N*_*m*_(0, *I*). In our model, since we aim to bridge the informative peak embedding *U* by using *Z*, we introduce an additional regularization term incorporating peak embeddings into the latent space. Specifically, we modify the latent factor *Z* as *Z*_*j*_ ∼ *Q*_Φ_(*z*_*j*_|*G*_*j*_, *u*_*j*_, *b*_*j*_). Therefore, we have

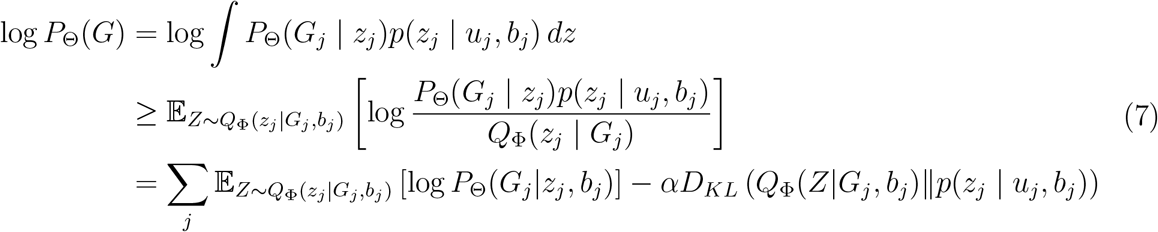

Now, we further decompose the KL divergence term *D*_*KL*_(*q*_*ϕ*_(*z* | *x*)∥*p*(*z* | *α, c*)). Given the conditional prior *p*(*z* | *α, c*), we can express it as:

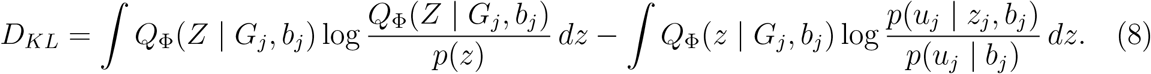

The first term simplifies to:

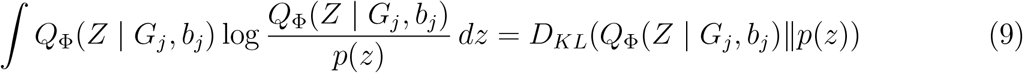

which represents the KL divergence between the variational distribution and the standard prior *p*(*z*). The second term can be rewritten as:

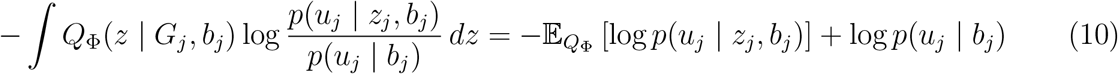

Ignoring the constant term log *p*(*u*_*j*_ | *b*_*j*_), the final form of the PEIVI loss function is:

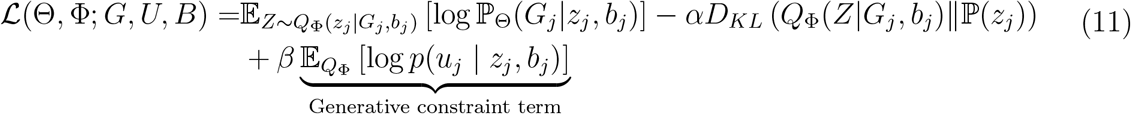

This result shows that *u*_*j*_ contributes an additional generative constraint term, which encourages the latent representation *z* to align with the peak embedding *u*_*j*_ given the condition *b*_*j*_. And since that *u*_*j*_ is fixed, this term can be approximated using a regularization function:

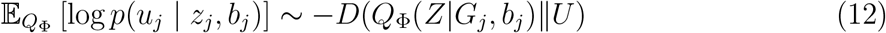

which leads to the final loss function ℒ(Θ, Φ; *G, U, B*):

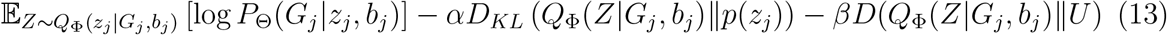

where *u*_*j*_ represents a deterministic precomputed peak embedding vector. This generative constraint term enforces alignment between the latent posterior *Q*_Φ_(*z*|*x*) and the domain-specific embedding *U*, which captures important peak-informed characteristics. To simplify model complexity while ensuring the optimization process is differentiable, we then approximate the divergence using the L2 distance metric (see Supplementary Note 1 for detailed approximation derivation of the distance function *D*(·)). Therefore, the revised objective function becomes:

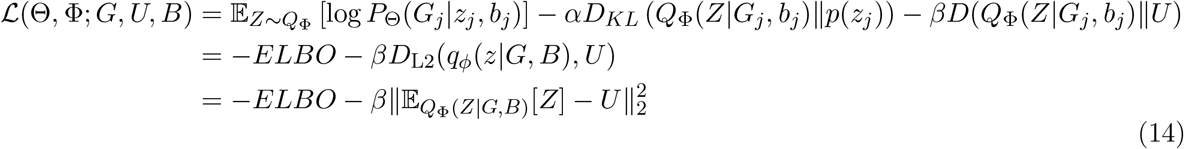

where *α >* 0 and *β >* 0 are hyperparameters that control the relative contribution of the embedding alignment term. Meanwhile, we also notice that both the genescore matrix and the incorporation of peak information into the bridge embedding will naturally introduce strong batch effects. To remove the batch effect from the integrated embedding, we apply maximum mean discrepancy (MMD) loss on *z*_*j*_. The MMD loss quantifies the extent of discrepancy between two distributions, and it was employed to align the latent cell embeddings across different batches^39^. As we described above, we denote the set of batches as *B* = *{b*_1_, *b*_2_, · · ·, *b*_*j*_*}*. Therefore, we have

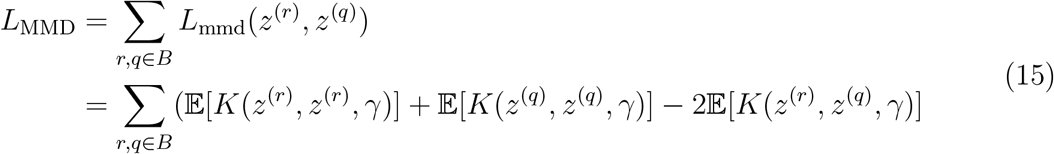

in which *K*(*·, ·, γ*) is a Gaussian kernel function that has 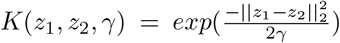, where *γ* is a hyperparameter.

By aggregating the loss function ℒ(Θ, Φ; *G, U, B*) and the MMD loss *L*_MMD_, we can formulate the ultimate optimization target of PEIVI

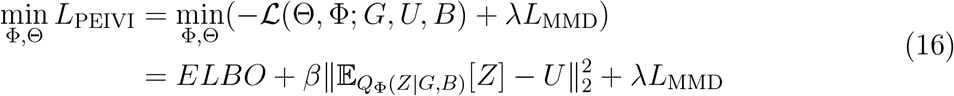

Complete pseudo-code of PEIVI is included in Supplementary Note 2. During training, EpiPack utilizes a staged training strategy to train PEIVI, that is, a pre-training stage is applied before MMD is used, in which the model learns peak embedding features, thus incorporating peak information into gene score space. Subsequently, MMD is applied with the regularization of the peak embedding to merge latent spaces from different reference sources or batches. Adam gradient descent is used to optimize the object loss function, and the reparametrization trick is employed to sample from the approximate posterior.

### Pretrained reference mapping

After training the model from scratch with reference datasets, we can conduct transfer learning to map the query data onto the reference latent embedding space; that is, fine-tuning the pre-trained model. More specifically, after integrating reference datasets *de novo*, pre-trained model *M* ^*ref*^ with parameters *θ*^*ref*^ and reference embedding *z*^*ref*^ can be saved and transferred to a newly initialized bridge VAE model *M* ^*query*^. Since the query batches are not located in the pre-trained model and can even contain different numbers of batches from different sources compared with the reference data, EpiPack will add new batch nodes with re-initialized weights to the first layer and replace the original query node. Weight initialization of the batch units follows Kaiming Initialization^62^. The parameters *θ*^*query*^ of the fine-tuned model can be expressed as

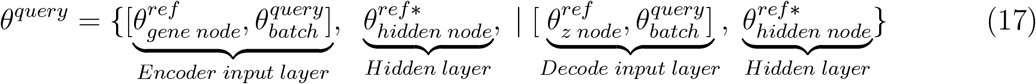

where *θ* represents fine-tuned weights and *θ*^∗^ represents fixed weight. In order to maximize the preservation of biological information within the mapping embedding space, EpiPack fixes all the hidden layers while fine-tuning the first layer of both the Encoder and Decoder.

The training procedure follows the same way as the EpiPack integration function. Batch-specific embeddings of the query datasets will be added to the uniform embedding space by optimizing the following loss function. Since the newly introduced peak embedding *u*^′^include direct batch effect to *z*^′^, we added an extra MMD loss between *z*^′^to the reference embedding *z*

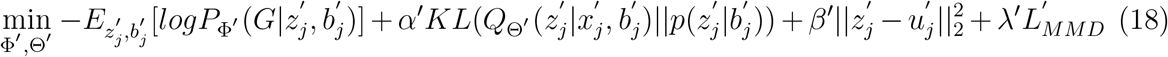

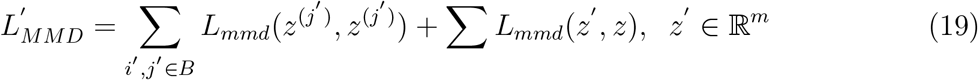

in which Φ^′^and Θ^′^are parameters of the query model with parameter set *θ*^*query*^.

### Classifier-based cell type annotation

Based on the integrated latent embeddings, the non-linear classifier is used to project the common space *L* to the classification space *C* and transfer the reference labels to the query dataset. Inspired by previous research, we realized that cell type imbalance can affect classifier performance. Therefore, we adopted the balanced sampler from scBalance^32^to construct training batches. Additionally, to achieve a more separable classification space that provides a better foundation for identifying new cell types, we constructed a prototype-based loss function to increase inter-class margins and decrease intra-class distances. We first initialize a prototype *c*_*m*_ for each cell type *m*, so for each cell type, we have distance 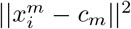, which represents intra-class compactness. Then, a normalized softmax is used to ensure inter-class sparsity. The total loss function is:

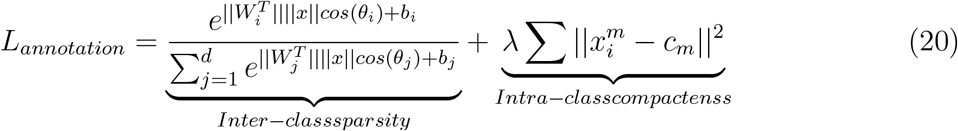

in which *x* means the input cell embedding. 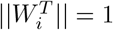 and *b* = 0. The hyperparameter *λ* controls the degree of centralization.

### Global OOR detection

Successful reference mapping and classification space projection provide a solid foundation for detecting OOR cell types. EpiPack global OOR detector subsequently employs predicted cell types in the classification space to detect potential new cell types.

The global detector first calculates the Mahalanobis distance based on the previous annotation result for each query cell. Let the random variable *K*, which is a dimension of the reference embedding generated by the EpiPack classifier, follow the Gaussian distribution *K* ∼ 𝒩(*µ, σ*^2^). Thus for each cell in the reference, we have a random vector 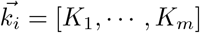 and a corresponding cell type **c**_*i*_ ∈ *{*1, · · ·, *C}*. Therefore, for each cell 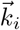, we have 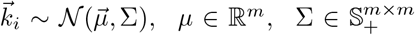 which is a multivariate Gaussian distribution. Considering the biological variance that exists in the cells, we assume that the multivariate normal distribution is anisotropic, which also corresponds with our observation that each cluster is not a standard circle. Thus, on the measurable reference space Ω in which 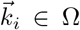, we can cell type-conditional distribution 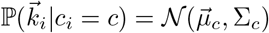 where

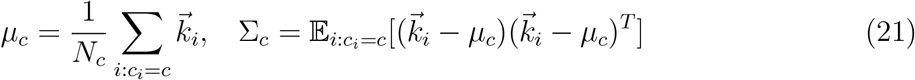

After that, we further evaluate whether the label assigned by the classifier is confident enough. Since we consider a discrete cell type, within-type density can be ignored, and we can assume a globally linear space for all those cell types. We have a conditional Gaussian. For each query cell 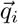 with a temporarily assigned label *c*_*i*_, we can measure the distance between 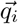 to the pre-defined distribution cell type-conditional distribution 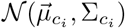 by

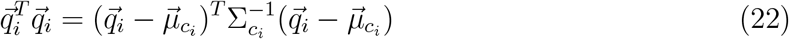

which is also the confidence ellipse of the class-conditional distribution of *c*_*i*_.

We then apply significance tests on the filtering result. Since manually setting the reject threshold is inappropriate in different annotation cases, here we implemented a significance-test-based method to discover OOR cells. We observed that the distance 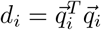 follows the chi-square distribution

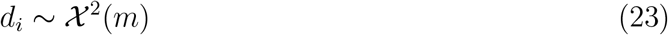

where *m* is the dimension of 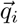. This can also be proved mathematically. Because the discrete OOR cell type is located out of the confidence region of the pre-defined population, though it is still assigned a false label by the classifier, we can thus discover such query cells by setting the minimum allowance p-value to reject the preliminary annotation. Normally, we recommend setting the threshold p-value to 1e-2 for the balance of the true positive rate and false discovery rate.

### Local OOR detection

The local OOR detector identifies subtle, cluster-specific state shifts by diffusing group information (control vs. perturbation) over a learned, anisotropic graph kernel. The method consists of four components: (i) joint graph construction, (ii) an edge-feature–driven learnable attention kernel *K*_*θ*_, (iii) bi-directional residual propagation (BRP), and (iv) node-level significance testing with FDR control.

#### Mutual kNN graph

Given the harmonized embedding *Z* ∈ ℝ^*n×d*^from PEIVI and a binary group vector *y* ∈ *{*0, 1*}*^*n*^(1=perturbation, 0=control), we first construct a mutual kNN graph on *Z* (typical *k* ∈ [15, 30], *k* ∈ **N**). Let *E* denote the set of directed edges (*i* → *j*) retained only if *i* is among *j*’s kNN and *j* is among *i*’s kNN to suppress spurious one-way links.

For each (*i, j*) ∈ ℰ we compute a low-dimensional feature vector *ϕ*_*ij*_ ∈ ℝ^*F*^designed to encode geometry and sampling context:

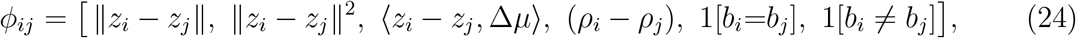

where Δ*µ* = mean(*Z*|*y*=1) − mean(*Z*|*y*=0) (robustly estimated), *ρ*_*k*_ is a nearest neighbor density proxy (mean of neighbor distances), and *b*_*i*_ denotes batch (if available). Features are standardized per dimension. The direction term ⟨*z*_*i*_ − *z*_*j*_, Δ*µ*⟩, which is an interproduct, promotes propagation along biologically plausible state axes; the density term *ρ*_*i*_ − *ρ*_*j*_ suppresses diffusion across low-density boundaries.

#### Learnable attention kernel *K*_*θ*_

To train the smooth kernel for the graph, we first compute a Gaussian kernel on existing edges as a geometry-only teacher,

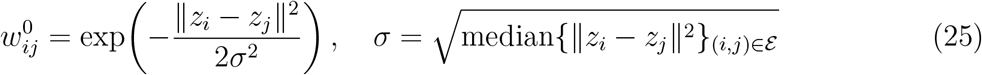

row-normalized to 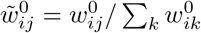. This provides a stable, isotropic baseline that anchors learning without using labels.

A small MLP maps edge features to non-negative scores *α*_*ij*_ = MLP_*θ*_(*ϕ*_*ij*_) and obtain a row-stochastic kernel by

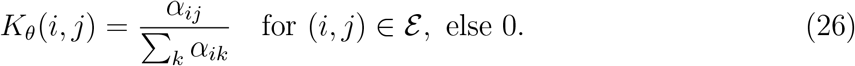

Next, training minimizes a composite loss over edges:

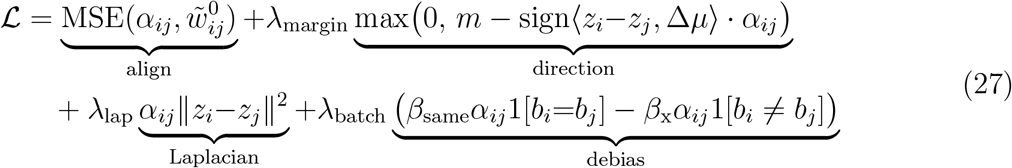

The align term keeps *K*_*θ*_ near a geometrically reasonable family; the direction and Laplacian terms encourage anisotropic yet local propagation; the batch regularizer penalizes within-batch preference and promotes cross-batch-consistent neighborhoods. By default, we use a 2 layer MLP (hidden 128, ReLU, dropout 0.1), Adam (lr = 10^−3^), margin *m* = 0.5.

#### Bi-directional residual propagation (BRP)

With the trained kernel, we diffuse query and reference signals separately with residual re-injection (to mitigate over-smoothing):

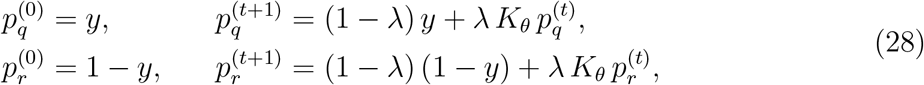

for *t* = 0, …, *T* − 1 with *λ* ∈ [0.85, 0.95] and *T* = 20−50 (or until ∥*p*^(*t*+1)^ − *p*^(*t*)^∥_1_*/n <* 10^−5^). We report per-cell log-odds and prior-corrected probabilities:

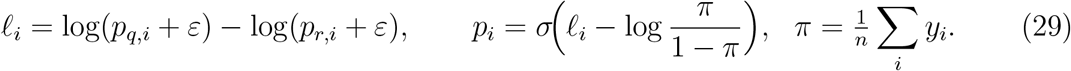

#### Significance testing and calling

We estimate a null for each node by permuting *y R* times (default *R* = 500) and re-running BRP to obtain 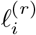. One-sided p-values are

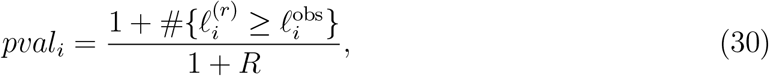

with Benjamini–Hochberg correction to *q*_*i*_. (Optionally, batch-stratified permutations preserve within-batch query fractions.) We call local OOR cells using FDR plus an effect threshold, e.g. *q*_*i*_ ≤ 0.05 & *p*_*i*_ ≥ 0.9.

#### Enrichment score

We calculate enrichment score *ES* for each cell by using the probability score *p*_*i*_

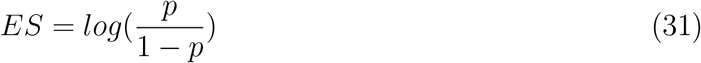

which represents the density of the specific experiment group.

### Hyper-parameters setting

Due to its deep learning architecture, EpiPack involves numerous hyperparameters. To optimize these settings, we conducted hyperparameter searches using the Hg38 PBMC benchmark dataset. Our exploration included algorithmic hyperparameters, such as the weight of the MMD loss (*λ*_*MMD*_) and the peak embedding regularization coefficient (*β*), as well as model hyperparameters, including the depth of the encoder and decoder, dropout rate, training batch size, and learning rate, among others. Detailed descriptions are provided in Extended Data Fig 3,4. The results indicate that our algorithm maintains robustness across a wide range of model hyperparameters. For algorithmic hyperparameters, we selected the set that yielded the best integration and mapping performance, which was then used to obtain the benchmark results presented in this study. Detailed tables of all model hyperparameters can be found in Supplementary Table 1-3.

### Pseudo-disease dataset simulation

To simulate partial-overlap shifts that mimic local differential abundance and the emergence of out-of-reference (OOR) states caused by perturbation, we applied a latent space editing framework to integrated scATAC-seq embeddings. Starting from an AnnData object containing a low-dimensional joint embedding *Z* ∈ ℝ^*n×d*^, annotated batch identities (with designated query/perturbed and reference/control subsets), and cell type labels, we selected a target population for perturbation. Within this population, we defined a one-dimensional perturbation axis *w* in the latent space, either by computing the centroid difference between query and reference subsets of the same type, or by extracting the first principal component from the query subset. The axis was normalized 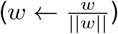, and each cell received a projection score 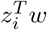. To induce a controlled partial overlap, we removed opposite tails of the score distributions from the query and reference (e.g., the lowest fraction in the query and the highest fraction in the reference, or vice versa), creating distributions that remain overlapping but misaligned along *w*. For benchmarking purposes, retained query cells at the extreme opposite tail to the removed fraction were annotated as OOR states, providing ground-truth labels for emerging shifted populations.

### Differential accessible peak analysis

We performed differential analysis of the COVID-associated OOR groups using Signac. First, we added DNA sequence motif information with the getMatrixSet and AddMotifs functions. We then applied the FindMarkers function with only.pos = FALSE and test.use = “LR” to identify peaks that were up- or down-regulated in OOR populations relative to the corresponding healthy control clusters. Subsequently, we used the FindMotifs function to retrieve significantly enriched motifs among the up-regulated peaks, and performed functional enrichment analysis of the associated genes using the enrichGO function.

### Identification of motif activities and differential analysis between OOR group and health control

We further performed fine-resolution differential motif analysis by computing motif scores for each cell using chromVAR. Specifically, we applied the RunChromVAR function in Signac, which calls the chromVAR package with the genome set to BSgenome.Hsapiens.UCSC.hg38. Differential motif activity was then assessed with the FindMarkers function, using mean.fxn = rowMeans and fc.name = “avg_diff” based on chromVAR z-scores. We set only.pos = TRUE to identify both up- and down-regulated motifs in the OOR groups. Finally, we visualized the results with ggplot2 by generating volcano plots of the differences in average TF accessibility.

### Datasets

#### Hg38 PBMC dataset

We curated the Hg38 PBMC dataset using publicly available 10x Genomics Human Health PBMC scATAC-seq data. This dataset comprises five samples, 38,853 cells, and ten distinct cell types. Before benchmarking, the data was subset to the 3,000 most variable genes. A detailed description of the dataset construction process is provided in Supplementary Note 2.

#### Cross reference genome PBMC dataset

Similarly, we constructed the cross-reference PBMC Dataset using publicly available 10x Genomics Human Health PBMC scATAC-seq data. This dataset consists of six samples, 42,652 cells, and ten cell types, with five samples derived from the Hg38 PBMC dataset and one sample generated from a 10x PBMC dataset based on the Hg19 reference genome. Before benchmarking, the data was subset to the 3,000 most variable genes. A detailed description of the dataset construction process is provided in Supplementary Note 2.

#### Small mouse cortex atlas

We constructed this dataset using two publicly available 10x Genomics Mouse Cortex scATAC-seq datasets. It consists of 12,445 cells spanning five cell types. Before benchmarking, the data was subset to the 3,000 most variable genes. A detailed description of the dataset construction process is provided in Supplementary Note 2.

#### Cardiac atlas

This dataset, published by Hocker et al.^2^, consists of 11 heart cell samples. We selected seven samples, comprising 31,308 cells and seven cell types, to evaluate the robustness of the model under different reference-to-query ratios. The gene score matrix was generated using snapATAC2. Before benchmarking, the data was subset to the 8,000 most variable genes. Ground-truth cell type labels were provided by the authors.

#### ccRCC dataset

This dataset, published by Kourtis et al.^43^, consists of early-stage clear cell renal cell carcinoma (ccRCC) scATAC-seq data. We selected seven samples, comprising 8,438 cells spanning ten cell types and states. The gene score matrix was generated using snapATAC2. Before the experiment, the data was subset to the 8,000 most variable genes. Ground-truth cell type labels were provided by the authors.

#### COVID dataset

In this dataset, we collected PBMC samples from eight healthy donors and one COVID-19 patient^46^. Among the healthy samples, five were obtained from the 10x PBMC dataset used in the benchmark experiments, and three were derived from donors of different ages. Two of these, together with the five 10x PBMC samples, were used to construct the reference atlas, while one healthy PBMC donor and the COVID-19 patient sample were designated as the control and query, respectively. The reference atlas dataset is first used to train the base model. Next, healthy and COVID samples are mapped to the reference space simultaneously by fine-tuning the pre-trained base model.

### Running methods

#### Reference mapping methods

We benchmarked the reference mapping performance of our model against various state-of-the-art methods, all of which rely on homogeneous feature spaces. These methods include:

- scArches scVI (v0.6.1): This method is based on the genescore matrix. We ran scArches scVI in Python by using the scArches package with the default parameters as suggested in the official tutorial. Based on the tutorial, 2000 HVGs are selected as the input of scArches scVI;
- PeakVI (v1.0.3): This method is based on the peak accessibility matrix. We ran PeakVI in Python by using the PeaVI package *load_query_data* function with the default parameters as suggested in the official tutorial;
- Seurat v3 (v5.2.0): This method is tested on both genescore and aligned peak matrices. We performed reference mapping following the provided tutorials and using default parameters.

#### De novo integration methods

We benchmarked the *de novo* data integration performance of our model against various state-of-the-art methods, all of which rely on homogeneous feature spaces. These methods include:

- LIGER (python v0.2.0): This method is based on the genescore matrix. We ran LIGER in Python by using the pyliger package (version 0.2.0) with the default parameters as suggested in the official tutorial. To align high-dimensional scATAC-seq datasets, we filtered the peaks that were not detected in more than 10% of the total cells. In the cross-reference genome integration test, the number of highly variable genes (HVGs) is set as 2000. HVGs are selected using Scanpy *highly*_*variable*_*genes* function;
- Harmony: We ran Harmony in R by using the harmony package (version 1.0) for the scATAC-seq peak matrix integration and ran the harmonypy package in Python (v 0.0.9) for gene score matrix integration. Default parameters as suggested in the official tutorial. The cell embedding used in Harmony came from Seurat LSI (peak embedding) and Scanpy PCA (genescore embedding);
- Combat (v1.4.5): This method is based on the peak accessibility matrix. Combat is a batch effect correction method developed for bulk data. We ran Combat in Python by using the Scanpy combat package;
- PeakVI (v1.0.3): This method is based on the peak accessibility matrix. We ran PeakVI in Python by using the scVI PEAKVI package with the default parameters as suggested in the official tutorial;
- Seurat CCA (Signac v1.10.0): We ran Seurat CCA in R by using Signac (version 1.10.0) *FindIntegrationAnchors* and *IntegrateEmbeddings* functions with the default parameters as suggested in the official tutorial. The method was trained based on the peak accessibility matrix;
- scVI (v1.0.3): We ran scVI in Python by using the scvi-tool package (version 1.0.3) with the default parameters as suggested in the official tutorial. Based on the tutorial, 2000 HVGs are selected as the input of scVI to integrate gene score matrices in the cross-reference-integration benchmarking experiment.

#### Cell label transfer methods

- Cellcano (v1.0.2): We ran Cellcano in the command line with the default parameters as suggested in the official tutorial;
- Seurat v3 (v5.2.0): we followed the tutorial and ran the model using default parameters;
- SVM: We used the SVC object from scikit-learn (v1.6.1) to train on the reference data and transfer cell labels. This method was trained based on the embedding space or genescore matrix.
- kNN: We used the KNeighborsClassifier object from scikit-learn (v1.6.1) to train on the reference data and transfer cell labels. This method was trained based on the embedding space

#### OOR detection methods

- kNN detector: Following the novel cell detection strategy implemented in scArches^12^, we constructed a kNN-based detector using the pynndescent package to assign uncertainty scores to each cell, where 0 indicates complete lack of confidence. We evaluated detectors with thresholds set at 0.5, 0.6, and 0.7, respectively.
- SVM detector: We implemented an SVC classifier using the scikit-learn SVM package, and obtained probabilistic confidence scores with the model.predict_proba function, where 0 indicates complete lack of confidence. Similar to the kNN detector, we tested thresholds of 0.5, 0.6, and 0.7.
- Milo (release 3.19): This method is based on neighborhood aggregation and employs negative binomial generalized linear models (NB-GLMs) to test for differential abundance (DA) of cells in single-cell datasets. We followed the official tutorial (https://github.com/MarioniLab/miloR) and ran Milo with default parameters.
- MELD (v1.0.2): This method is based on graph-based kernel density estimation, which generalizes kernel density estimation (KDE) from a regular spatial domain to a manifold represented by a cell–cell similarity graph, followed by label signal smoothing. We applied MELD following the official tutorial (https://github.com/KrishnaswamyLab/MELD/tree/main) and ran the method with default parameters.

### Evaluation metrics

In our study, the evaluation metrics can be divided into three categories: reference mapping metrics, cell label transfer metrics, and OOR detection metrics.

#### Metrics for the reference mapping task

- ARI: Adjusted rand index (ARI) score measured the extent to which cells of different types are correctly clustered, regardless of the batch used in the latent embedding. After clustering cells using latent embeddings obtained from different integration and reference mapping methods, we calculated the ARI by comparing the clustering labels with the ground-truth cell labels, which is defined as:

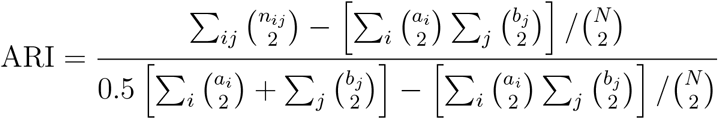

where *n*_*ij*_ represents the number of samples in the intersection of the *i*-th true cluster and *j*-th predicted cluster, *a*_*i*_ and *b*_*j*_ are the respective cluster sizes, and *N* is the total number of samples. The clustering labels were generated using the Leiden clustering algorithm.
- NMI: Normalized mutual information (NMI) score was also used to measure the extent to which cells of different types are correctly clustered using latent embeddings, which is defined as:

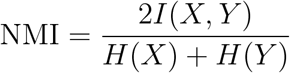

where *I*(*X, Y*) is the mutual information between the ground truth labels *X* and predicted clusters *Y*, and *H*(*X*), *H*(*Y*) are their respective entropies. NMI ranges from 0 to 1, with higher values indicating better clustering consistency. We used NMI to compare cell type labels with Leiden clusters calculated on the integrated dataset.
- Cell type ASW: Cell type ASW (average silhouette width) was used to measure the relationship between the intra-cluster distance of cells and the inter-cluster distance to the nearest cluster. This is also a cell-type-level metric. As defined by a recent benchmarking paper^18^,

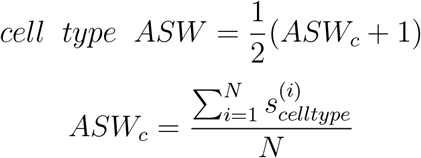

in which 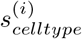 is the silhouette width for the *i*th cell, and N is the total number of cells.
- Batch ASW: Batch ASW (average silhouette width) is a variant of cell type ASW, used to measure the effectiveness of batch mixing. Unlike cell type ASW, batch ASW calculates the silhouette width using the batch labels of each cell *i*.

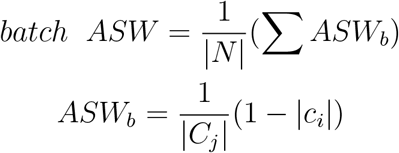

in which *C*_*j*_ is a cell set with the label *j*. |*N*| is the number of the set of unique cell labels.
- iLISI: Integration Local Inverse Simpson’s Index (iLISI) score quantifies batch mixing within local neighbor-hoods. A higher iLISI value indicates effective batch mixing, whereas a lower iLISI suggests poor integration with residual batch structure. iLISI is computed for each cell based on the Inverse Simpson’s Index (ISI), which measures the diversity of batch labels among its nearest neighbors. Given a dataset of *C* cells, iLISI for cell *i* is defined as:

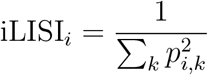

where *p*_*i,k*_ represents the proportion of cells from batch *k* within the k-nearest neighbor (k-NN) graph. The overall dataset-level iLISI score is obtained by averaging across all cells:

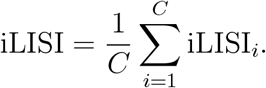
- kBET: k-Nearest-Neighbor batch estimation (kBET) algorithm assesses whether the label composition within a cell’s k-nearest neighbors is similar to the expected global label composition. The method evaluates batch mixing by performing a 𝒳^2^goodness-of-fit test for each cell, comparing the observed batch distribution within its local neighborhood to the expected distribution under perfect batch mixing. Given a dataset of *C* cells, kBET computes the rejection rate, defined as the fraction of cells where the null hypothesis (uniform batch distribution) is rejected. The final kBET score is given by:

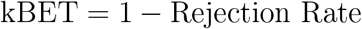

where values close to 1 indicate successful batch mixing, and values near 0 suggest the presence of residual batch effects.

##### Biological conservation

In our experiments, we used adjusted rand index, normalized mutual information, and cell type ASW to measure the biological conservation of batch integration and reference mapping. According to the recently published scIB benchmark package^18^, we averaged these three metrics to summarize them as a single biological conservation metric.

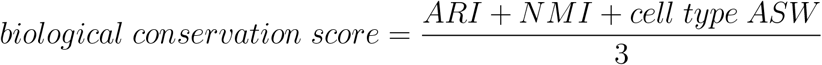

##### Batch correction

Similarly, in our experiments, we used iLISI, k-Nearest-Neighbor batch estimation, and batch ASW to measure the batch correction of batch integration and reference mapping. According to the recently published scIB benchmark package, we averaged these three metrics to summarize them as a single batch correction metric.

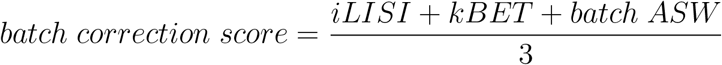

##### Overall score

According to the recently published scIB benchmark package, the overall score is calculated by.

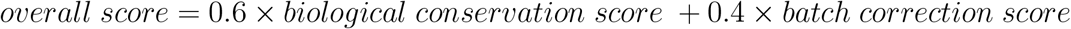

#### Metrics for the cell label transfer task

For the cell label transfer task, we used the Macro F1 score and Weighted F1 score to assess predictive performance. The F1 score for a given cell type *c* is defined as:

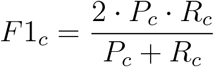

where 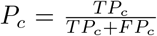 (precision) and 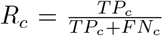 (recall) are computed per class, with *T P*_*c*_, *FP*_*c*_, and *FN*_*c*_ denoting the true positives, false positives, and false negatives for class *c*, respectively.

- Macro F1: The Macro F1 score computes the unweighted mean across all *C* cell types:

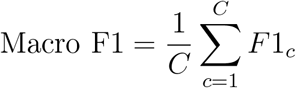

ensuring equal contribution from each class, regardless of class size.
- Weighted F1: The Weighted F1 score accounts for class imbalance by weighting each class’s F1-score by its relative support *w*_*c*_ (i.e., proportion of cells in that cell type):

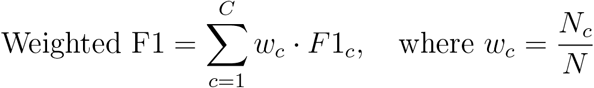

where *N*_*c*_ is the number of cells in class *c* and *N* is the total number of cells.

#### Metrics for the OOR detection task

For global and local OOR detection tasks, we used the True Positive Rate (TPR) and False Discovery Rate (FDR) to assess predictive performance.

- True Positive Rate (TPR). The TPR score measures the proportion of true out-of-reference (OOR) cells that are correctly identified as OOR. It is defined as:

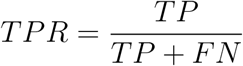

where *T P* denotes the number of true positives (correctly identified OOR cells), and *FN* denotes the number of false negatives (missed OOR cells). A higher TPR indicates better sensitivity in detecting OOR cells.
- False Discovery Rate (FDR). The FDR score measures the proportion of predicted OOR cells that are actually false positives (i.e., reference cells incorrectly classified as OOR). It is defined as:

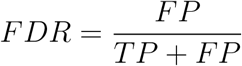

where *FP* denotes the number of false positives (reference cells incorrectly classified as OOR). A lower FDR indicates higher precision in OOR detection.

## Data availability

All datasets used in this paper are previously published and freely available. The Mouse cortex datasets and Hg38 and Hg19 PBMC datasets are downloaded from the 10X Genomics dataset portal, for which detailed URLs are provided in Supplementary Note 2. The Cardiac atlas dataset is available at Gene Expression Omnibus under accession number GSE165837. Corresponding peak files and metadata can be downloaded at http://ns104190.ip-147-135-44.us/CARE_portal/ATAC_data_and_download.html. The clear cell renal cell carcinoma (ccRCC) dataset is available at Gene Expression Omnibus under accession code GSE181064. The COVID dataset used in the case study is available at Gene Expression Omnibus under accession code GSE173590.

## Code availability

The EpiPack python package is available at https://github.com/ZhangLabGT/EpiPack. Tutorials of the package are also available at https://epipack.readthedocs.io/en/main/.

## Author contribution

Y.C. and X.Z. conceived this study. Y.C. designed and implemented the models. Y.C. developed the package and carried out the evaluation and data analysis. Y.C. and X.Z. wrote the paper. X.Z. supervised the work.

## Acknowledgements

This work was supported by National Institutes of Health grant R35GM143070.

## Competing interests

The authors declare no competing interests.

---

**Extended Data Figure 1.**
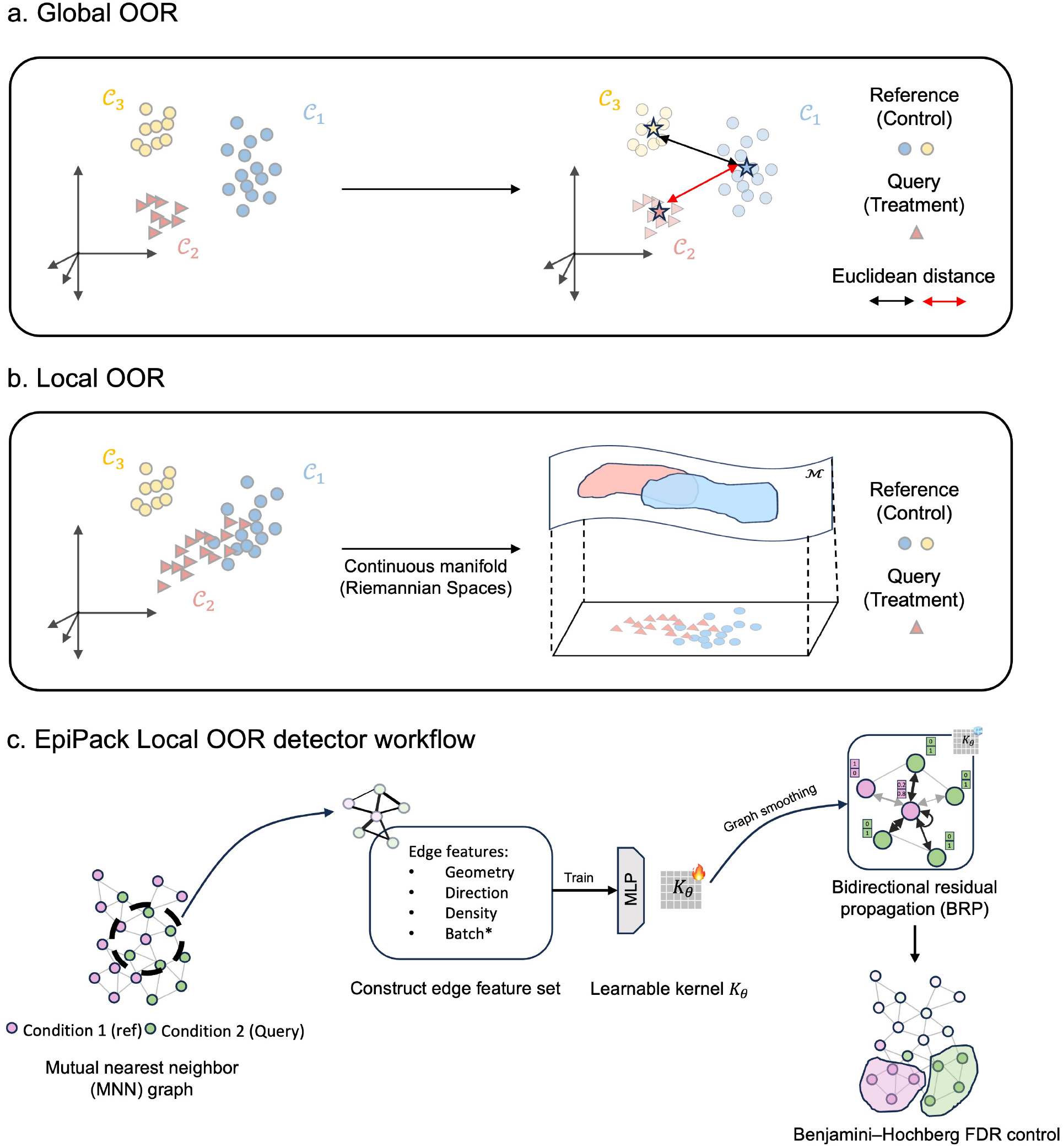
Schematic of the Global-Local OOR framework in EpiPack. **a. Global OOR**: OOR cell types that are biologically distinct from in-reference types (e.g., CD4 T vs. B cells) form isolated clusters in the joint latent embedding space. **b. Local OOR**: Local OOR represents subtle, continuous perturbations of known cell types or states, often arising from gradual transitions (e.g., cell activation, dysfunction, or differentiation). Unlike global OOR, these states do not form isolated clusters but instead lie along smooth, continuous Riemannian manifolds in the latent space, where the global Euclidean metric is no longer valid. **c. Local OOR detector**. The detector constructs a mutual nearest neighbor graph between reference and query cells, augments edges with geometry, direction, density, and batch features, and trains an MLP to learn an adaptive kernel *K*_*θ*_. This kernel guides bidirectional residual propagation to capture subtle local deviations while avoiding oversmoothing. Significant local OOR regions are identified through Benjamini–Hochberg FDR control.

**Extended Data Figure 2.**
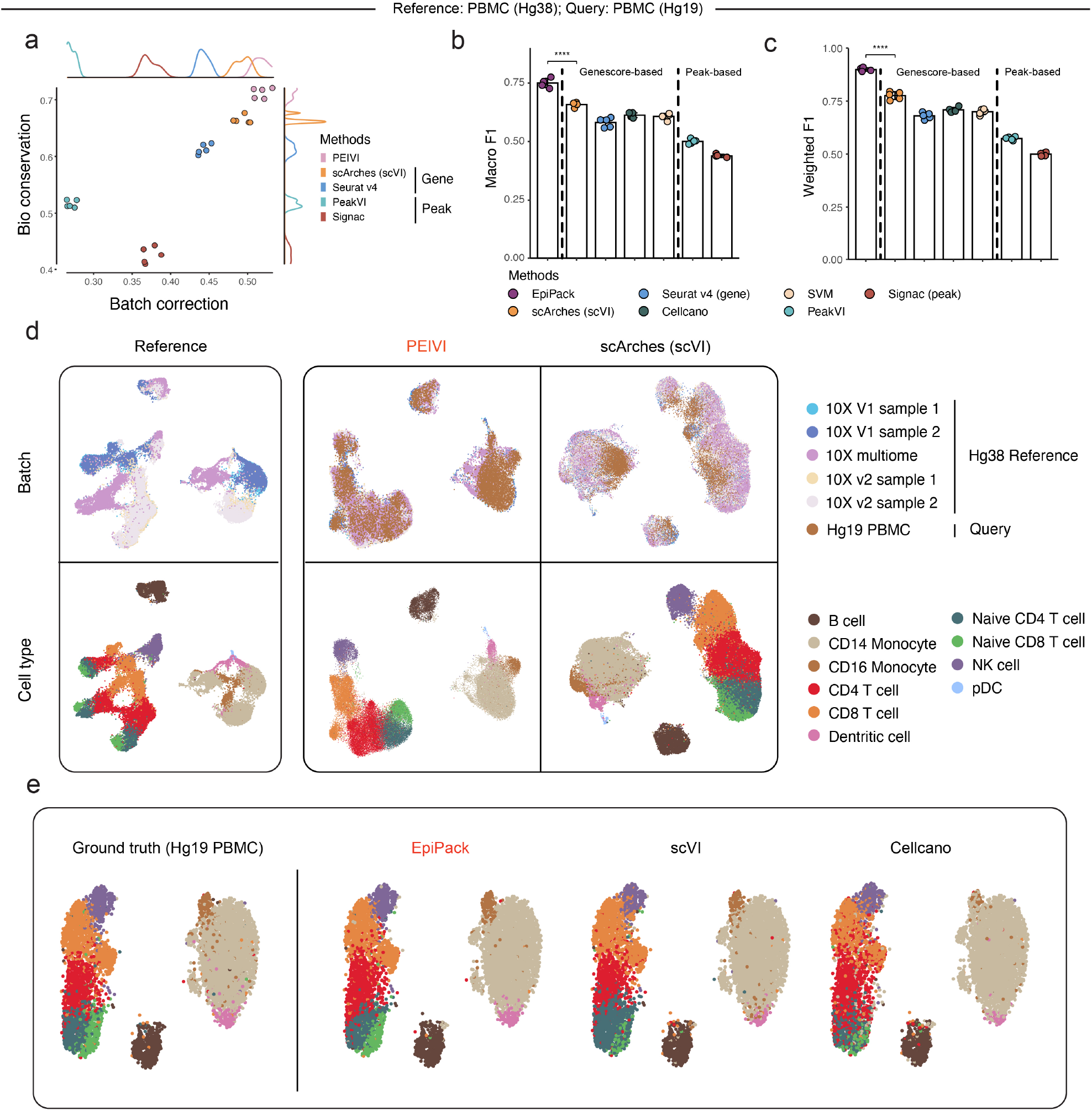
Incorporating heterogeneous features substantially improves scATAC-seq mapping and annotation across reference genomes. **a**. Overall scores for the benchmarked models’ biological conservation and batch correction performance on the cross-reference genome setting using five PBMC datasets aligned to the Hg38 genome as the reference and one PBMC dataset aligned to Hg19 as the query. **b-c**. Comparison of cell type annotation accuracy across methods using macro F1 (b) and weighted F1 (c) scores. Scores are averaged over five repeated experiments; error bars denote standard deviation. Asterisks indicate statistical significance (two-sided t-tests, **** means p < 0.0001). **d**. UMAP visualization of the joint embedding spaces learned by PEIVI and scArches. **e**. Cell label transfer results on the Hg19 PBMC dataset. Compared to scVI and Cellcano, PEIVI demonstrates markedly higher accuracy, especially for rare populations such as naive CD8 T cells, dendritic cells, and CD16 monocytes.

**Extended Data Figure 3.**
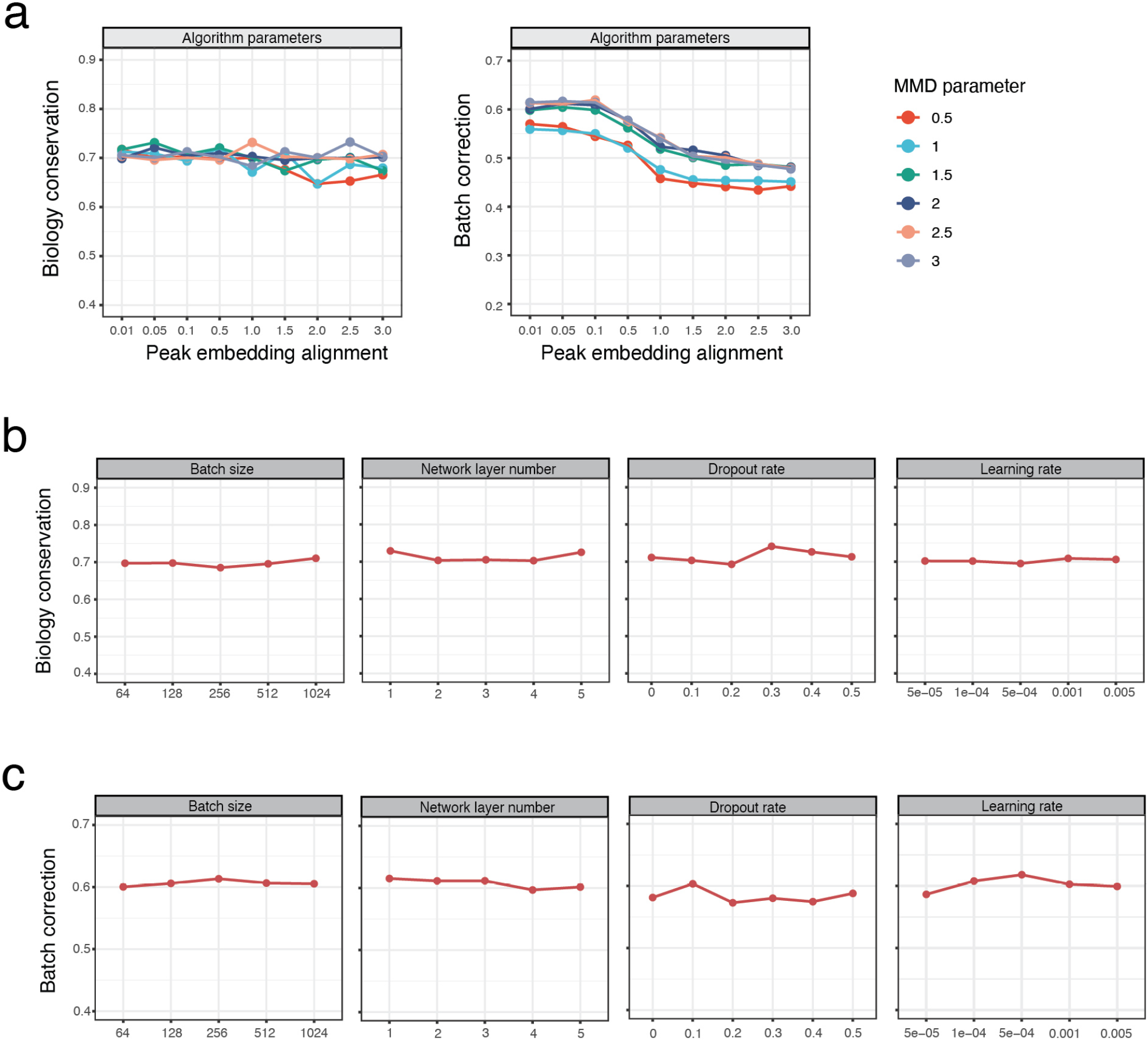
Hyperparameter sensitivity analysis for PEIVI in de novo reference construction. **a**. Grid search over two key algorithmic hyperparameters: the MMD regularization weight (colored lines) and the peak embedding constraint coefficient (x-axis), evaluated on biological conservation (left) and batch correction (right) scores. PEIVI maintains stable integration performance across a wide range of settings. **b-c**. Robustness analysis of model-level hyperparameters, assessed by biological conservation (b) and batch correction (c) scores. “Batch size” is the number of training samples in each mini-batch during optimization. “Network layer number” is the number of hidden layers in the encoder and decoder networks. “Dropout rate” is the fraction of units randomly dropped during training. “Learning rate” is the step size used for gradient descent updates.

**Extended Data Figure 4.**
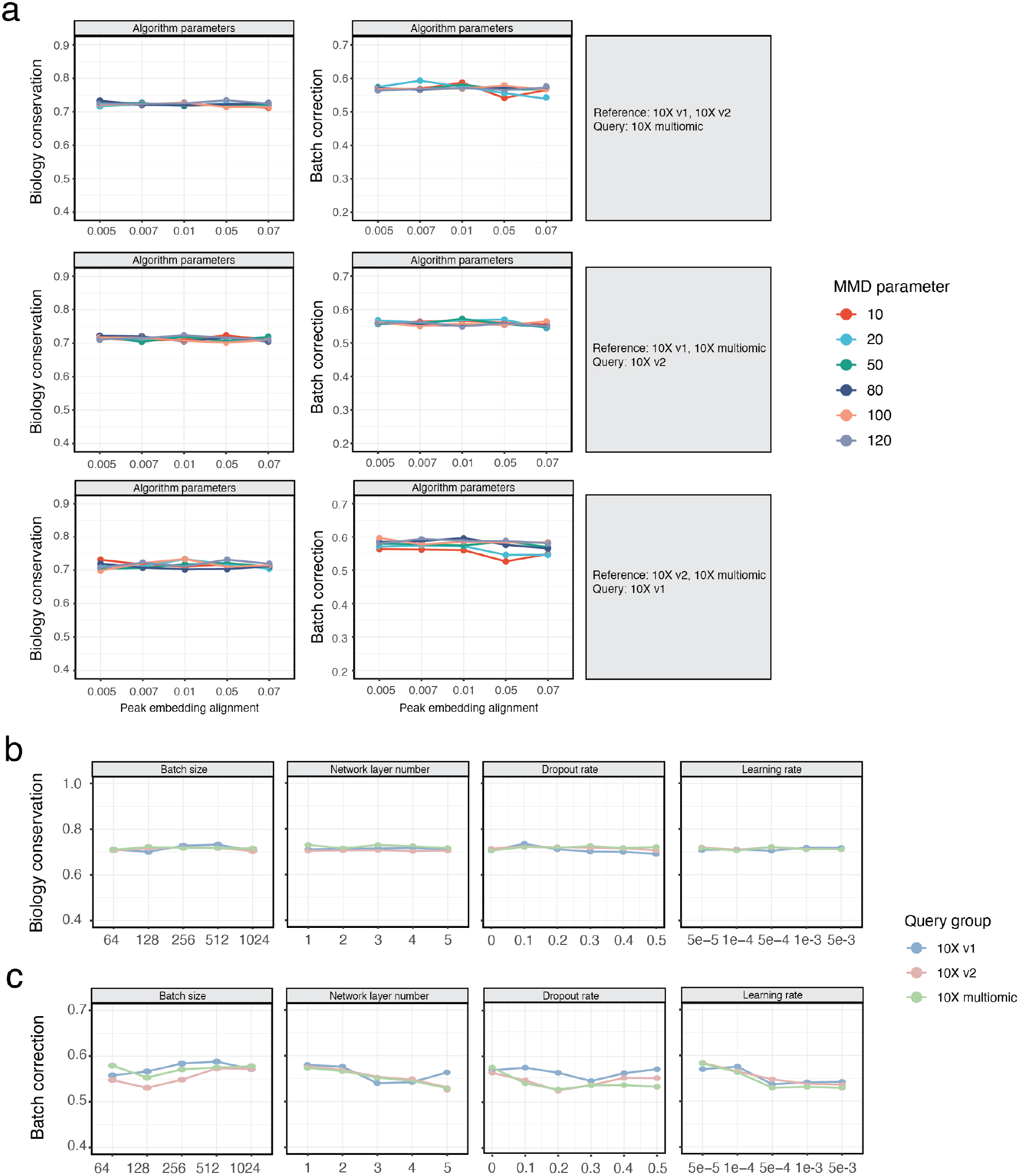
Hyperparameter robustness analysis of PEIVI for scATAC-seq query mapping tasks. **a**. Grid search over two core algorithmic hyperparameters: the MMD regularization coefficient (colors) and the peak embedding constraint coefficient (x-axis), across three reference-query configurations. Performance is evaluated using biological conservation (left) and batch correction (right) scores. PEIVI maintains stable performances across a wide range of hyperparameter combinations. **b-c**. Model-level hyperparameter robustness evaluation across the same three query groups, with lines color-coded by query condition.

## Notes

### Competing Interest Statement

The authors have declared no competing interest.

