## Supplementary table note and figures for "Discovering cell types and states from reference atlases with heterogeneous single-cell ATAC-seq features"

#### Supplementary Figure 9

Supplementary Fig 1 - Overall scores for the benchmarked models' biological conservation and batch correction performance 9
Supplementary Fig 2 - UMAP visualization of reference mapping results. 10
Supplementary Fig 3 - - Cell label transfer performance on the joint embedding space across methods to reflect nearest neighbor structure preservation in unsupervised reference mapping. 11
Supplementary Fig 4 - Weighted F1 and Macro F1 scores of the benchmarked models in the supervised cell label transfer setting 11
Supplementary Fig 5 - Data integration benchmarking results. 12
Supplementary Fig 6. UMAP visualization of de novo integration results on mouse cortex scATAC-seq data. 13
Supplementary Fig 7. UMAP visualization of de novo integration results on human PBMC scATAC-seq data. 13
Supplementary Fig 8. UMAP visualizations of PEIVI mapping at different ref:query ratios (human PBMC). 14
Supplementary Fig 9 - UMAP visualization of EpiPack's cluster separability on rare cell populations at different proportions 15
Supplementary Fig 10. UMAP visualization of the predicted labels against the ground-truth label on the query 2 experiment set. 16
Supplementary Fig 11. Boxplots of TPR and FDR benchmarking results with CD8 T cells as the out-of-reference (OOR) population. 16
Supplementary Fig 12. UMAP visualization of the gene score highlights in reference atlas embedding. 17
Supplementary Fig 13. UMAP visualization of the gene score highlights in the health-COVID joint embedding. 17
Supplementary Fig 14. Differential peak analysis in the B cell between the OOR group and the health control cluster. 18

#### References 19

### 1 Supplementary Table

#### 1.1 Supplementary table 1: PEIVI architecture and training hyperparameters

| Name | Layer name | Dimension | Dropout | Normalization | Activation |
|---|---|---|---|---|---|
| **Inputs** | | | | | |
| data | – | Num of Genes | | | – |
| conditions | – | Num of Batches | | | – |
| conditions | – | Dim of peak embedding | | | – |
| **Encoder** | | | | | |
| Layer_1 | FC | 256 | 0.1 | BN | LeakyReLU |
| Layer_2 | FC | 256 | 0.1 | BN | LeakyReLU |
| mean | FC | 50 | | | Linear |
| var | FC | 50 | | | Linear |
| **Decoder** | | | | | |
| Layer_1 | FC | 256 | 0.1 | BN | LeakyReLU |
| Layer_2 | FC | 256 | 0.1 | BN | LeakyReLU |
| mean | FC | Num of Genes | | | ReLU |
| theta | FC | Num of Genes | | | Softplus |
| **Hyperparameters** | | | | | |
| Loss | NB | | | | |
| Optimizer | Adam | | | | |
| Learning Rate | 0.0001 | | | | |
| Weight Decay | 5e-4 | | | | |
| Batch Size | 512 | | | | |
| clamp | 0.01 | | | | |
| training stage 1 | 50 epoch | | | | |
| training stage 2 | 100 epoch | | | | |

**Table 1.** PEIVI architecture and training hyperparameters.

#### 1.2 Supplementary table 2: Peak encoder architecture and training hyperparameters.

| Name | Layer name | Dimension | Dropout | Normalization | Activation |
|---|---|---|---|---|---|
| **Inputs** | | | | | |
| data | – | Num of Peaks | | | – |
| **Encoder** | | | | | |
| Layer_1 | FC | 256 | 0.1 | LN | LeakyReLU |
| Layer_2 | FC | 256 | 0.1 | LN | LeakyReLU |
| latent | FC | 50 | | | Linear |
| **Decoder** | | | | | |
| Layer_1 | FC | 256 | 0.1 | LN | LeakyReLU |
| Layer_2 | FC | 256 | 0.1 | LN | LeakyReLU |
| rate | FC | Num of Peaks | | | Sigmoid |
| **Hyperparameters** | | | | | |
| Loss | BCE (with lib size) | | | | |
| Optimizer | Adam | | | | |
| Learning Rate | 0.0001 | | | | |
| Weight Decay | 5e-4 | | | | |
| Batch Size | 512 | | | | |

**Table 2.** Peak encoder architecture and training hyperparameters.

#### 1.3 Supplementary table 3: Epipack classifier architecture and training hyperparameters.

| Name | Layer name | Dimension | Dropout | Normalization | Activation |
|---|---|---|---|---|---|
| **Inputs** | | | | | |
| data | – | Dim of joint embedding | | | – |
| **Encoder** | | | | | |
| Layer_1 | FC | 64 | 0.1 | BN | ReLU |
| latent | FC | 30 | | | Linear |
| **Hyperparameters** | | | | | |
| Loss | Combined loss | | | | |
| Optimizer | Adam | | | | |
| Learning Rate | 0.0001 | | | | |
| Weight Decay | 5e-4 | | | | |
| Batch Size | 128 | | | | |

**Table 3.** Classifier architecture and training hyperparameters.

#### 2 Supplementary Note

##### 2.1 Supplementary Note 1: Heterogeneous transfer constraint term approximation

As mentioned in the Methods section, the loss function of PEIVI can be derived from the Evidence Lower Bound (ELBO) as follows:

$$\begin{aligned}
\log \mathbb{P}_\Theta(G) &= \log \int \mathbb{P}_\Theta(G_j \mid z_j) p(z_j \mid u_j, b_j)\, dz \\
&\geq \mathbb{E}_{Z\sim Q_\Phi(z_j|G_j,b_j)} \left[ \log \frac{\mathbb{P}_\Theta(G_j \mid z_j)\mathbb{P}(z_j \mid u_j, b_j)}{Q_\Phi(z_j \mid G_j)} \right] \\
&= \sum_j \mathbb{E}_{Z\sim Q_\Phi(z_j|G_j,b_j)} \left[ \log \mathbb{P}_\Theta(G_j|z_j,b_j) \right] - \alpha D_{KL}\left( Q_\Phi(Z|G_j,b_j) \| \mathbb{P}(z_j \mid u_j, b_j) \right) \\
&= \sum_j \mathbb{E}_{Z\sim Q_\Phi(z_j|G_j,b_j)} \left[ \log \mathbb{P}_\Theta(G_j|z_j,b_j) \right] - \underbrace{\alpha D_{KL}\left( Q_\Phi(Z|G_j,b_j) \| \mathbb{P}(z_j) \right)}_{\text{KL term}} - \underbrace{\beta D(Q_\Phi(Z|G_j,b_j) \| U)}_{\text{Generative constraint term}}
\end{aligned} \tag{1}$$

where $u_j$ represents a deterministic precomputed peak embedding vector. Among these terms, the prior vector $u_i$ and its constraint on the latent embedding, referred to as the "Generative constraint term", play a crucial role in enabling heterogeneous transfer in the latent space. Therefore, the key challenge lies in optimizing this loss function, specifically in determining the appropriate metric $D(Q_\Phi(Z|G_j,b_j)\|U)$ to measure this distance.

And because the anchor $U \in \mathbb{R}^{d_z}$ is a deterministic latent code, treating it as a Dirac measure $\delta_{U_i}$ provides a principled notion of proximity between the posterior and the anchor via the 2-Wasserstein distance[1]:

$$W_2^2\big(Q_\Phi(Z \mid G_i, b_i),\, \delta_U\big) \;=\; \mathbb{E}_{Z\sim Q_\Phi}\big[\|Z-U\|_2^2\big].$$

This identity shows that the squared Euclidean distance between a posterior sample and $U_i$ is a *stochastic, unbiased estimator* of an optimal-transport distance to the point anchor[2,3]. Concretely, drawing a reparameterized sample $z_i = \mu_i + \sigma_i \odot \varepsilon$ with $\varepsilon \sim \mathcal{N}(0, I)$ yields

$$\widehat{D}_i \;=\; \|z_j - u_j\|_2^2, \qquad \mathbb{E}\big[\widehat{D}_i\big] = W_2^2\big(Q_\Phi, \delta_{u_j}\big),$$

and averaging $L$ samples $\frac{1}{L}\sum_{\ell=1}^{L} \|z_i^{(\ell)} - u_j\|_2^2$ reduces estimator variance if desired. In contrast, the KL divergence to a Dirac target is ill-posed (infinite for any posterior with nonzero variance), so the Wasserstein route both avoids degeneracy and endows the constraint with a true metric geometry.

Operationally, adding the penalty $\beta\|z_j - u_j\|_2^2$ to the objective injects scATAC-derived information into the latent code by exerting a sample-level pull toward $U$. The gradient $\partial\|z_j - u_j\|_2^2/\partial z_j = 2(z_j - u_j)$ propagates through the reparameterization to the encoder parameters Φ, encouraging draws $z_j \sim Q_\Phi(Z \mid G_j, b_j)$ to concentrate near $U$ while the standard ELBO terms preserve data reconstruction and prior regularization. Thus,

$$D\big(Q_\Phi(Z \mid G_j, b_j) \,\|\, U\big) \;\approx\; \|z_j - u_j\|_2^2$$

is not a heuristic, but the Monte-Carlo instantiation of $W_2^2\big(Q_\Phi, \delta_U\big)$, providing a simple, differentiable, and theoretically grounded mechanism to fuse the anchor $U$ into the latent space.

#### 2.2 Supplementary Note 2: Pseudo-code of PEIVI

**Algorithm 1** PEIVI mappable latent space construction (two-stage training)

**Input:** scATAC-seq batches $\{b_j\}_{j=1}^{J}$; stage lengths $E_1 = 50$, $E_2 = 100$

**for** $j = 1$ to $J$ **do**
  Compute gene activity score matrix $G_j$ for $b_j$
**end for**
Obtain shared gene matrix $G = \bigcap_{j=1}^{J}\{G_j\}$
**for** $j = 1$ to $J$ **do**
  Initialize batch-specific autoencoder $u_j$ with parameters $\omega_{u_j}$; train independently
  **while** not converged (or until max epoch) **do**
    $\omega_{u_j} \leftarrow \arg\min\ L = \mathrm{BCELoss}(b_j, \tilde{b}_j)$
  **end while**
  Return batch-specific latent embedding $u_j$
**end for**
Initialize bridge autoencoder with parameters $\Phi, \Theta$
**Stage 1 (warm-up, $E_1 = 50$ epochs): optimize** $L_{\text{elbo}} + D$
**for** $e = 1$ to $E_1$ **do**
  Compute standard ELBO loss $L_{\text{elbo}}$ on $\{z_j\}$
  Compute alignment regularizer $D = \sum_{j=1}^{J}\sum_i \left\| z_j^{(i)} - u_j^{(i)} \right\|_2^2$
  $(\Phi, \Theta) \leftarrow \arg\min\ \big(L_{\text{elbo}} + D\big)$
**end for**
**Stage 2 (regularized integration, $E_2 = 100$ epochs): optimize** $L_{\text{elbo}} + D + L_{\text{MMD}}$
**for** $e = E_1 + 1$ to $E_1 + E_2$ **do**
  Compute $L_{\text{elbo}}$ on $\{z_j\}$
  Compute $D = \sum_{j=1}^{J}\sum_i \left\| z_j^{(i)} - u_j^{(i)} \right\|_2^2$
  Compute batch-wise MMD loss $L_{\text{MMD}}$ between $\{z_j\}$
  $(\Phi, \Theta) \leftarrow \arg\min\ \big(L_{\text{elbo}} + D + L_{\text{MMD}}\big)$
**end for**
Return integrated latent embedding $z\{z_j\}_{j=1}^{J}$

**Output:** Integrated embedding $z$, pre-trained model with parameter set $\theta^{ref} = (\Phi, \Theta)$

#### 2.3 Supplementary Note 3: Datasets preprocessing

##### (1). Small mouse scATAC-seq datasets (2 batches)

The mouse scATAC-seq batch integration dataset includes two scATAC-seq datasets. For these two datasets, we retrieved peak count matrix and fragment files from the 10X Genomics data portal. The batch 1 dataset is collected from "8k Adult Mouse Cortex Cells, ATAC v2, Chromium Controller" (https://www.10xgenomics.com/datasets/8k-adult-mouse-cortex-cells-atac-v2-chromium-controller-2-standard).

The batch 2 dataset is collected from "8k Adult Mouse Cortex Cells, ATAC v2, Chromium X" (https://www.10xgenomics.com/resources/datasets/8k-adult-mouse-cortex-cells-atac-v2-chromium-x-2-standard).

Both datasets are aligned on the GRCm38 (mm10) reference genome. We first preprocessed both datasets by filtering low quality cells and peak regions according to the standard tutorial of Signac. Then the gene score matrix is calculated for each dataset by the Signac *GeneActivity* function with default parameters (2kb upstream of the TSS), and the datasets were concatenated. As the peaks called per dataset differ, we used the union of the two peak files to obtain the merged peak dataset. Finally, we obtain a union peak matrix with two batches (12445 cells, 194403 peaks), a union gene score matrix with two batches (12445 cells, 3000 highly variable genes are selected), and two filtered peak count matrices.

We annotated the cells in five major cell types using typical marker genes in Signac (Astrocytes: Aldh1l1, Gfap, Gja1, S100b; Excitatory neurons: Satb2, Slc17a7, Slc17a8; Inhibitory neurons: Gad1, Gad2, Grik1; Microglia: Cd68, Cd14, Fcgr1, S100A8, S100A9; Oligodendrocytes: Mag, Mog, Olig1)

##### (2). Large human PBMC scATAC-seq datasets (5 batches)

The human PBMC scATAC-seq batch integration dataset includes five scATAC-seq datasets. For these five datasets, we retrieved the peak count matrix and fragment files from the 10X Genomics data portal. The batch 1 dataset is collected from "5k PBMCs from a Healthy Donor (Next GEM v1.1)" (https://www.10xgenomics.com/resources/datasets/5-k-peripheral-blood-mononuclear-cells-pbm-cs-from-a-healthy-donor-next-gem-v-1-1-1-1-standard-1-2-0).

The batch 2 dataset is collected from "10k PBMCs from a Healthy Donor (Next GEM v1.1)" (https://www.10xgenomics.com/resources/datasets/10-k-periphe-ral-blood-mononuclear-cells-pbm-cs-from-a-healthy-donor-next-gem-v-1-1-1-1-standard-2-0-0).

The batch 3 dataset is collected from "10k Human PBMCs, ATAC v2, Chromium Controller" (https://www.10x-genomics.com/resources/datasets/10khuman-pbmcs-atac-v2-chromium-controller-2-standard).

The batch 4 dataset is collected from "10k Human PBMCs, ATAC v1.1, Chromium X" (https://www.10xgenomics.com-/resources/datasets/10k-human-pbmcs-atac-v1-1-chromium-x-1-1-standard).

The batch 5 dataset is collected from "10k Human PBMCs, Multiome v1.0, Chromium X" (https://www.10xgenomics.com/resourc-es/datasets/10-k-human-pbm-cs-multiome-v-1-0-chromium-x-1-standard-2-0-0).

All these five datasets are aligned on the GRCh38 (Hg38) reference genome. We first preprocessed all datasets by filtering low quality cells and peak regions according to the standard tutorial of Signac[4]. Then the gene score matrix is calculated for each dataset by the Signac *GeneActivity* function with default parameters (2kb upstream of the TSS), and the datasets were concatenated. For de novo integration experiments, we used the union of the peak files to obtain the merged peak dataset. For peak-dependent reference mapping tools, the query data were filtered using the peak set of the reference model to ensure a consistent feature space. Finally, we obtain a union peak matrix with five batches, a union gene score matrix with five batches (3000 highly variable genes are selected), and five filtered peak count matrices.

Since the raw 10X PBMC datasets are not labeled, we manually annotated the cells in ten major cell types using typical marker genes in Signac (CD14 Monocyte: CD14, LYZ; CD16 Monocyte: FCGR3A, MS4A7; B cell: MS4A1, CD74; CD4 T cell: CD3, CD4, IL7R, S100A4; CD8 T cell: CD8A, NKG7; Dendritic cell: FCER1A, CST3; NK cell: NKG7, GNLY; Naive CD4 T cell: CD4, LEF1, CCR7; Naive CD8 T cell: CD8A, LEF1, CCR7; pDC: CD45R, BST2)

**(3). Cross reference genome human PBMC scATAC-seq datasets**

The cross reference genome scATAC-seq batch integration dataset includes 2 scATAC-seq datasets. One is based on the Hg38 reference genome and the other is based on the Hg19 reference genome. The Hg38 dataset is the multiomic 10X PBMC dataset we used in the Large human PBMC scATAC-seq dataset integration task ("10k Human PBMCs, Multiome v1.0, Chromium X"). The Hg19 dataset is collected from the "10k Peripheral Blood Mononuclear Cells (PBMCs) from a Healthy Donor" (`https://www.10xgenomics.com/resources/datasets/10-k-peripheral-blood-mononuclear-cells-pbm-cs-from-a-healthy-donor-1-standard-1-0-1`).

The cross reference genome scATAC-seq reference mapping dataset includes 6 scATAC-seq datasets. The reference set is based on the Hg38 reference genome (5 datasets, same as the Human Hg38 PBMC). And the query set is based on the Hg19 reference genome.

We first preprocessed all datasets by filtering low quality cells and peak regions according to the standard tutorial of Signac. Then the gene score matrix is calculated for each dataset by the Signac *GeneActivity* function with default parameters (2kb upstream of the TSS), and the datasets were concatenated. Due to the substantial variability of peak features between the hg38 and hg19 genome builds, we first performed peak region correction using liftOver[5]. The chain file for converting hg38 to hg19 (hg38ToHg19.over.chain.gz) is available from the UCSC Genome Browser file server (https://hgdownload.cse.ucsc.edu/goldenpath/hg38/liftOver/). After correction, we constructed a merged peak dataset by taking the union of the peak files across datasets for integration. For peak-dependent reference mapping tools, the query data were filtered using the peak set of the reference model to ensure a consistent feature space.

#### 3 Supplementary Figure

###### 3.0.1 Supplementary Fig 1 - Overall scores for the benchmarked models' biological conservation and batch correction performance

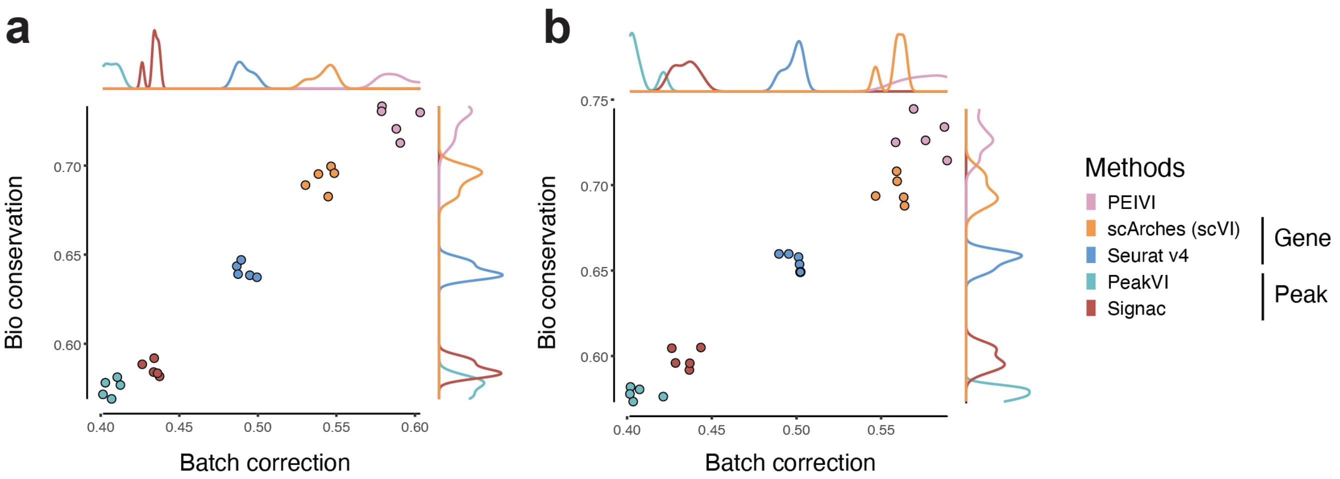

**Supplementary Fig. 1. a.** Benchmarking result for the experiment group "reference: 10x v2 and multiomics, query: 10x v1" (n=5 for 5 repeating experiments) **b.** Benchmarking result for the experiment group "reference: 10x v1 and v2, query: 10x multiomics" (n=5 for 5 repeating experiments)

##### 3.0.2 Supplementary Fig 2 - UMAP visualization of reference mapping results.

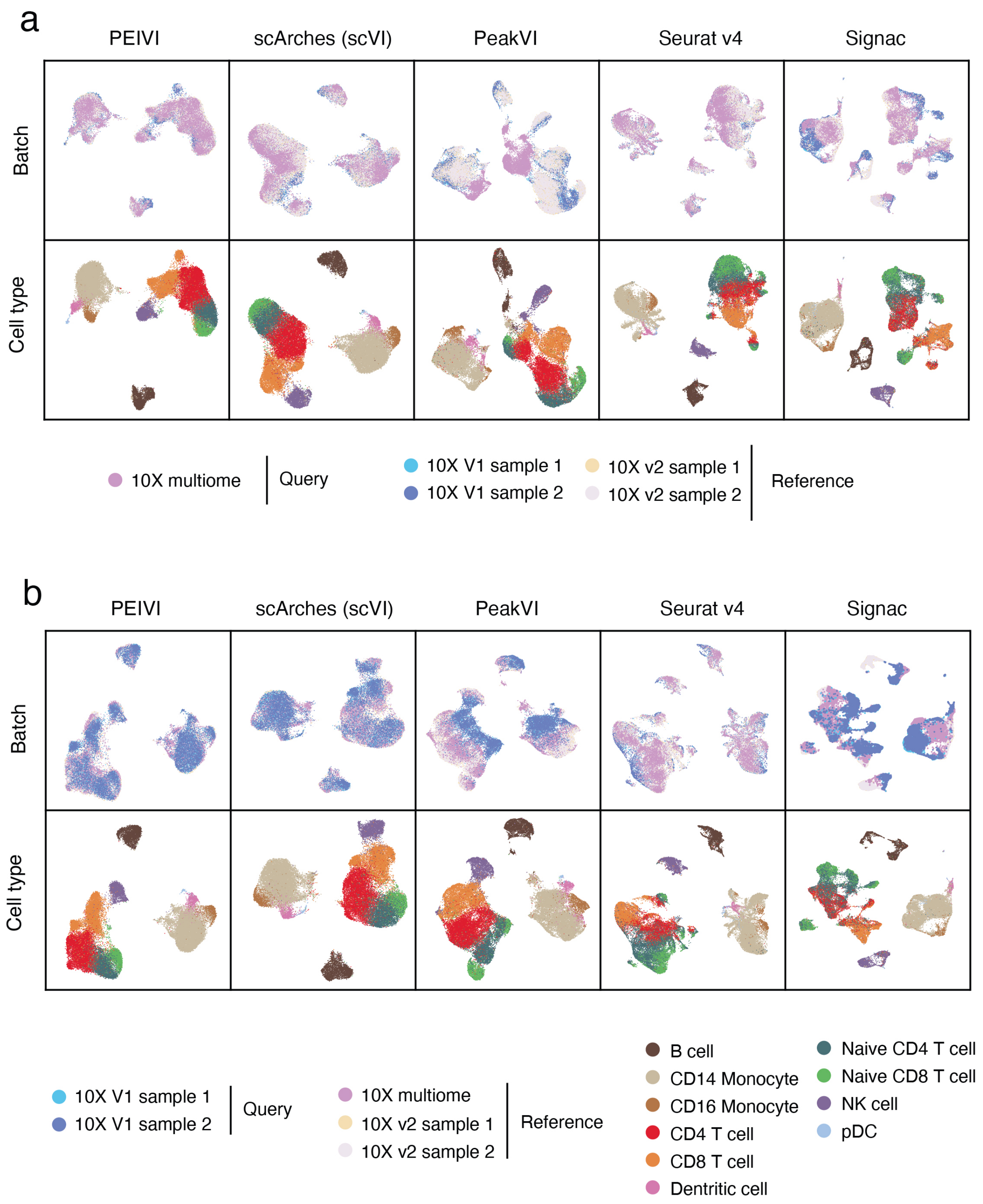

**Supplementary Fig. 2. a.** UMAP visualization result for the experiment group "reference: 10x v2 and multiomics, query: 10x v1" (n=5 for 5 repeating experiments) **b.** UMAP visualization result for the experiment group "reference: 10x v1 and v2, query: 10x multiomics" (n=5 for 5 repeating experiments)

##### 3.0.3 Supplementary Fig 3 - - Cell label transfer performance on the joint embedding space across methods to reflect nearest neighbor structure preservation in unsupervised reference mapping.

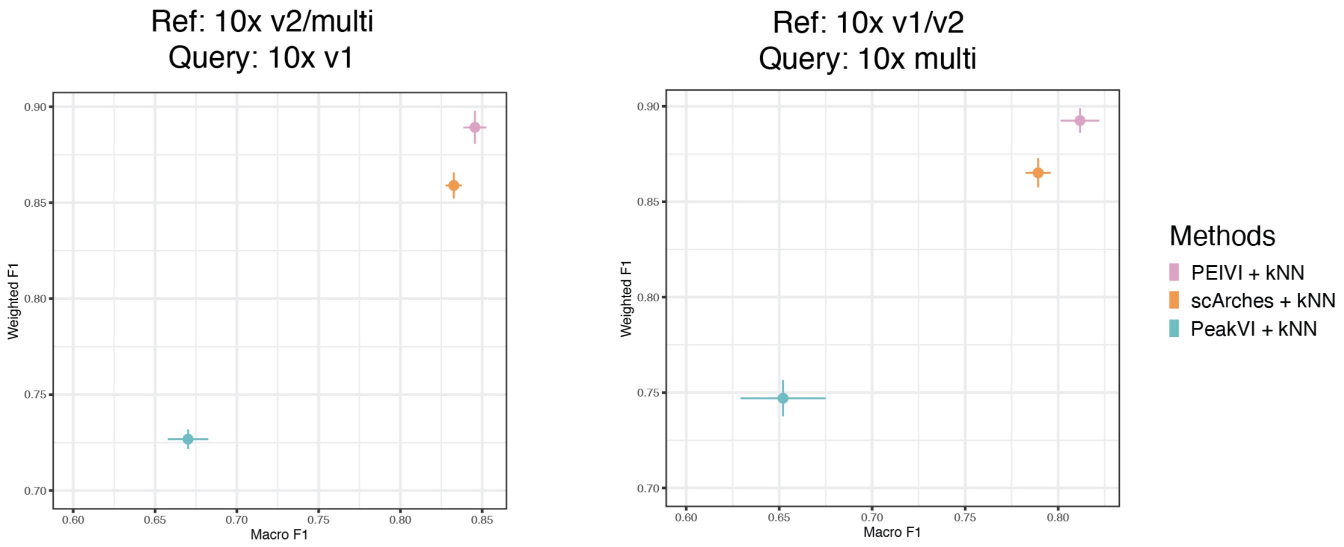

**Supplementary Fig. 3. a.** Benchmarking result for the experiment group "reference: 10x v2 and multiomics, query: 10x v1" (n=5 for 5 repeating experiments) **b.** Benchmarking result for the experiment group "reference: 10x v1 and v2, query: 10x multiomics" (n=5 for 5 repeating experiments)

##### 3.0.4 Supplementary Fig 4 - Weighted F1 and Macro F1 scores of the benchmarked models in the supervised cell label transfer setting

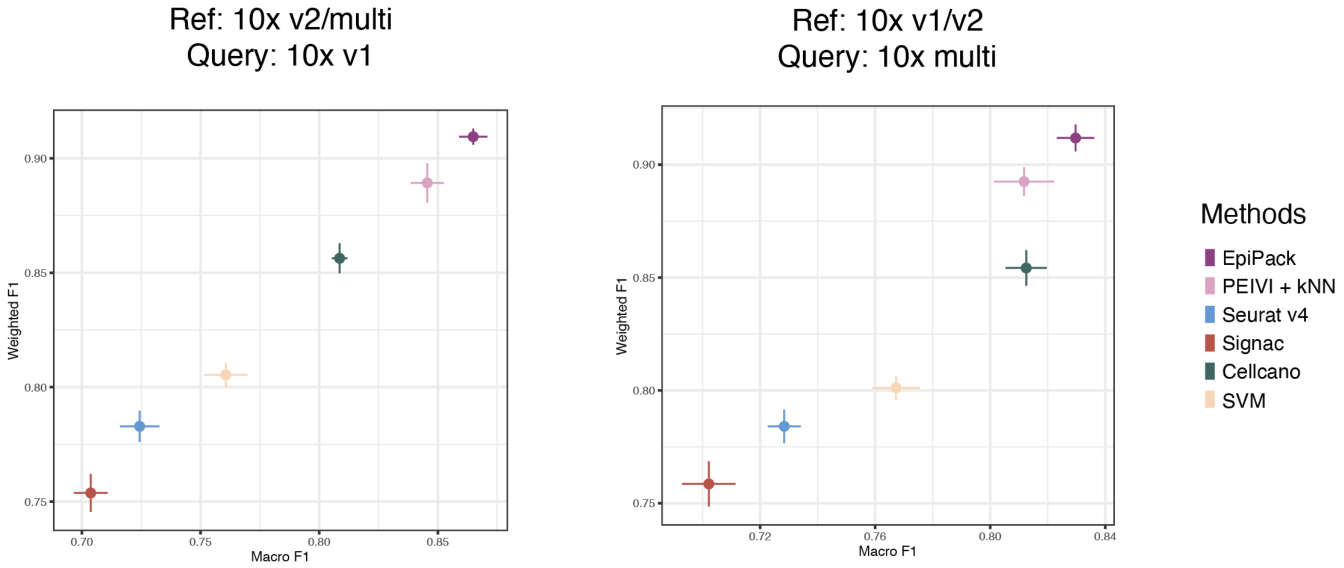

**Supplementary Fig. 4. a.** Benchmarking result for the experiment group "reference: 10x v2 and multiomics, query: 10x v1" (n=5 for 5 repeating experiments) **b.** Benchmarking result for the experiment group "reference: 10x v1 and v2, query: 10x multiomics" (n=5 for 5 repeating experiments)

##### 3.0.5 Supplementary Fig 5 - Data integration benchmarking results.

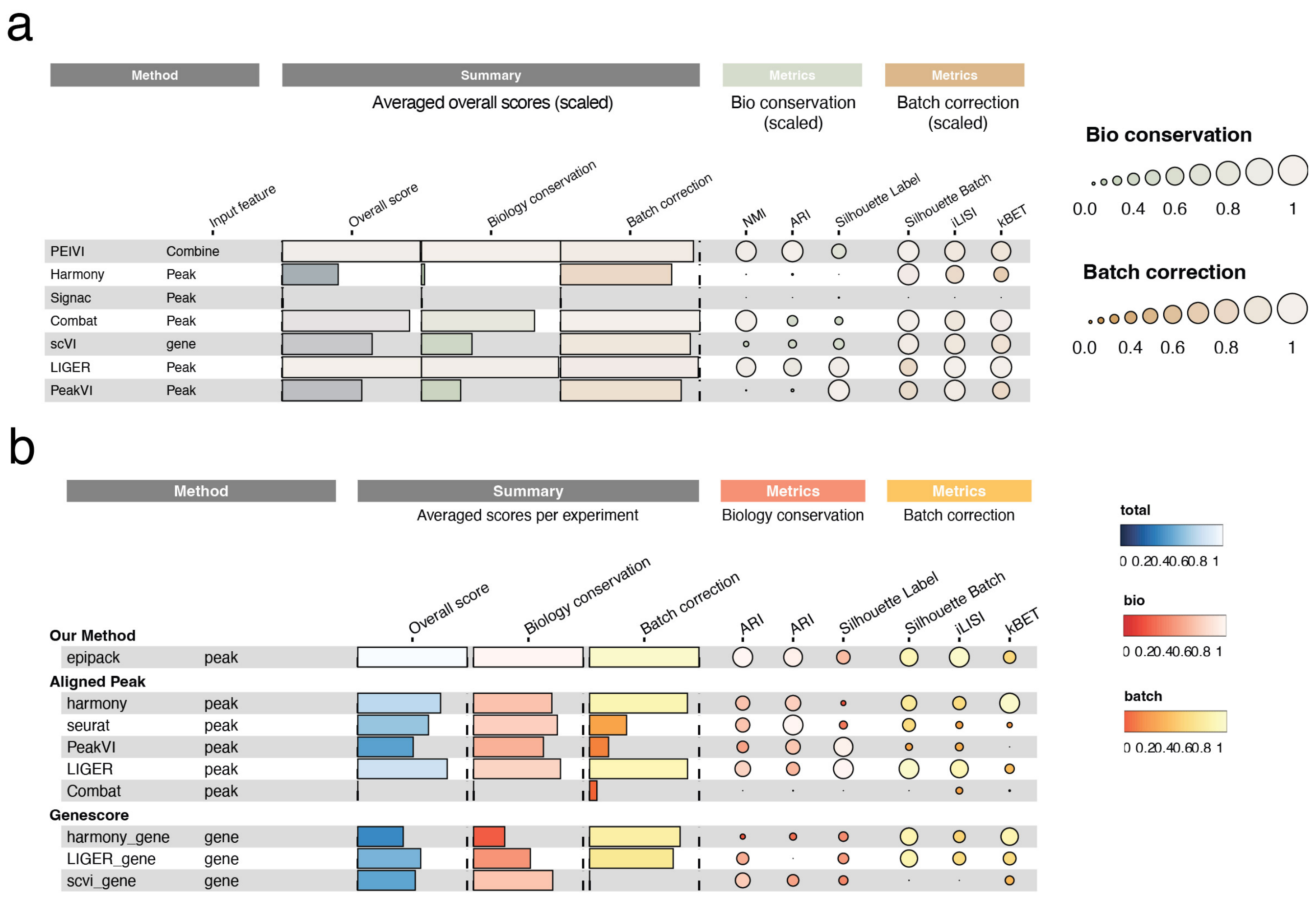

**Supplementary Fig. 5. a.** Benchmarking of PEIVI against six widely used data integration methods with different types of aligned input features on the mouse brain dataset. **b.** Benchmarking of PEIVI against six widely used data integration methods with different types of aligned input features on the cross reference genome setting.

##### 3.0.6 Supplementary Fig 6. UMAP visualization of de novo integration results on mouse cortex scATAC-seq data.

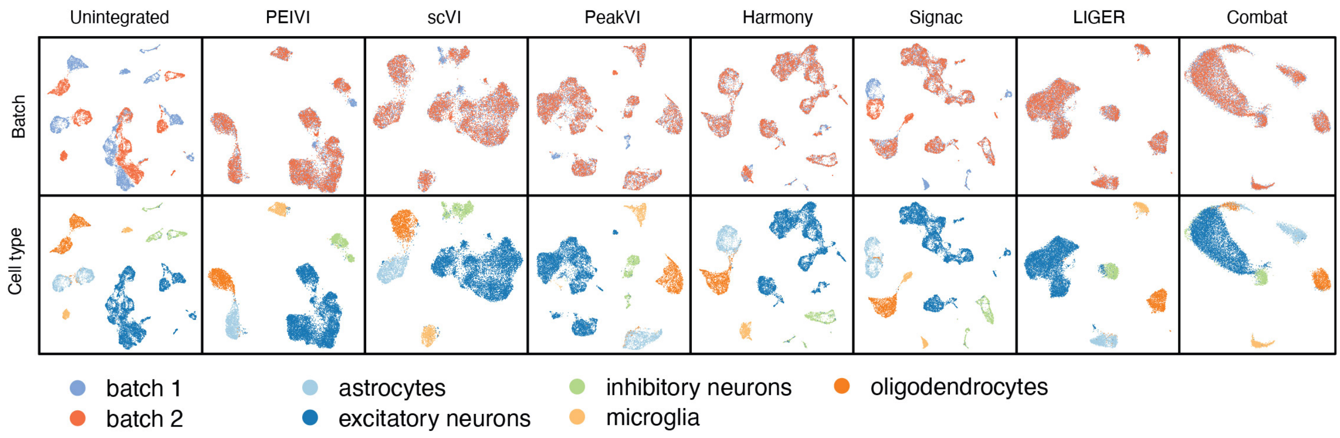

**Supplementary Fig. 6.** UMAP visualization of de novo integration results on mouse cortex scATAC-seq data.

##### 3.0.7 Supplementary Fig 7. UMAP visualization of de novo integration results on human PBMC scATAC-seq data.

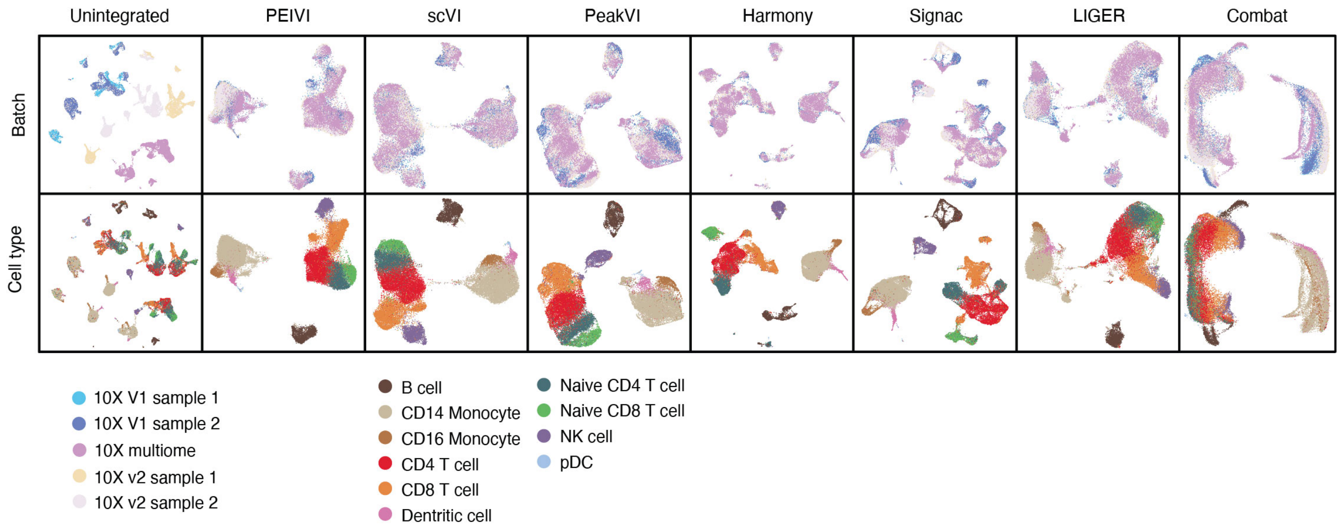

**Supplementary Fig. 7.** UMAP visualization of de novo integration results on human PBMC scATAC-seq data.

##### 3.0.8 Supplementary Fig 8. UMAP visualizations of PEIVI mapping at different ref:query ratios (human PBMC).

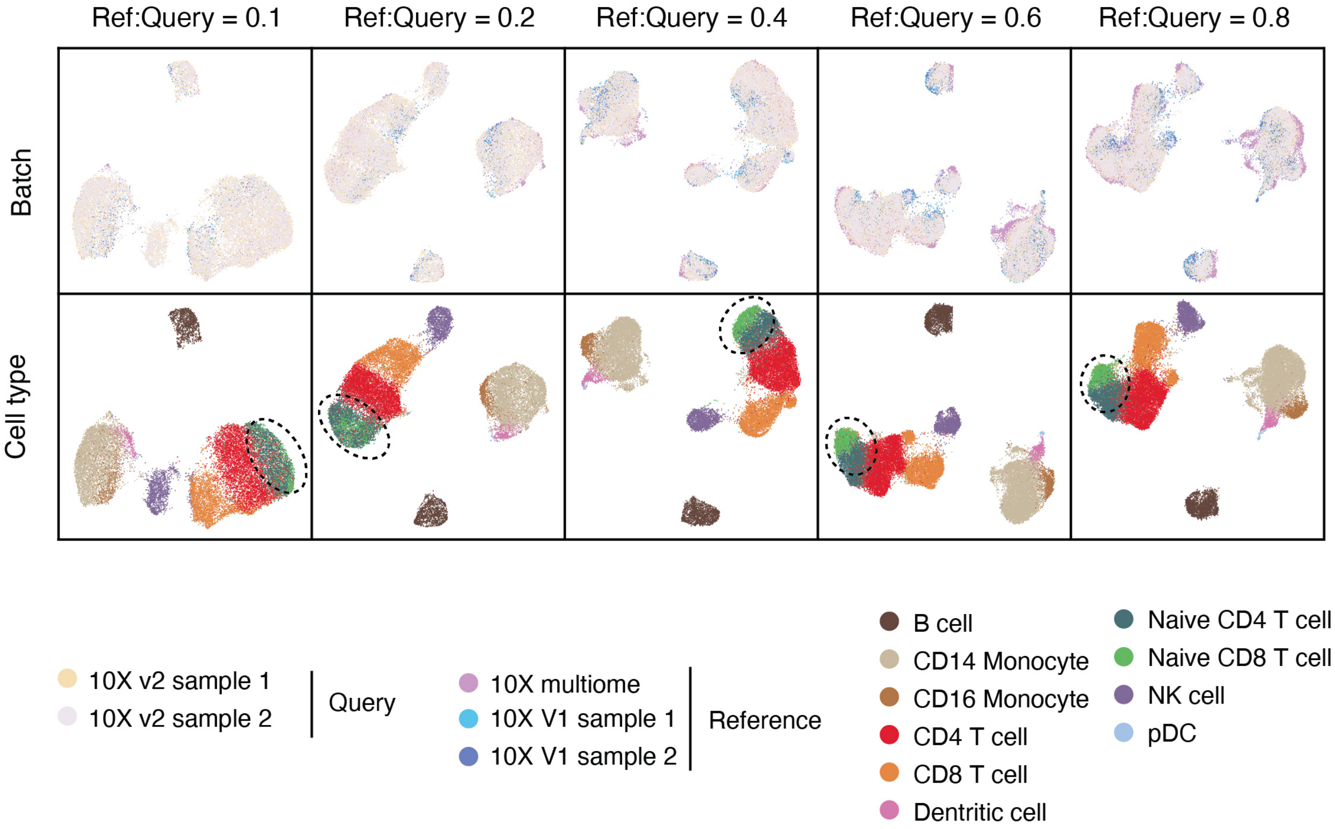

**Supplementary Fig. 8.** UMAP visualizations of PEIVI mapping at different ref:query ratios (human PBMC). Distinct Naive CD8 T cell clusters (highlighted by black circles) emerge when the ref:query ratio exceeds 0.6.

##### 3.0.9 Supplementary Fig 9 - UMAP visualization of EpiPack's cluster separability on rare cell populations at different proportions

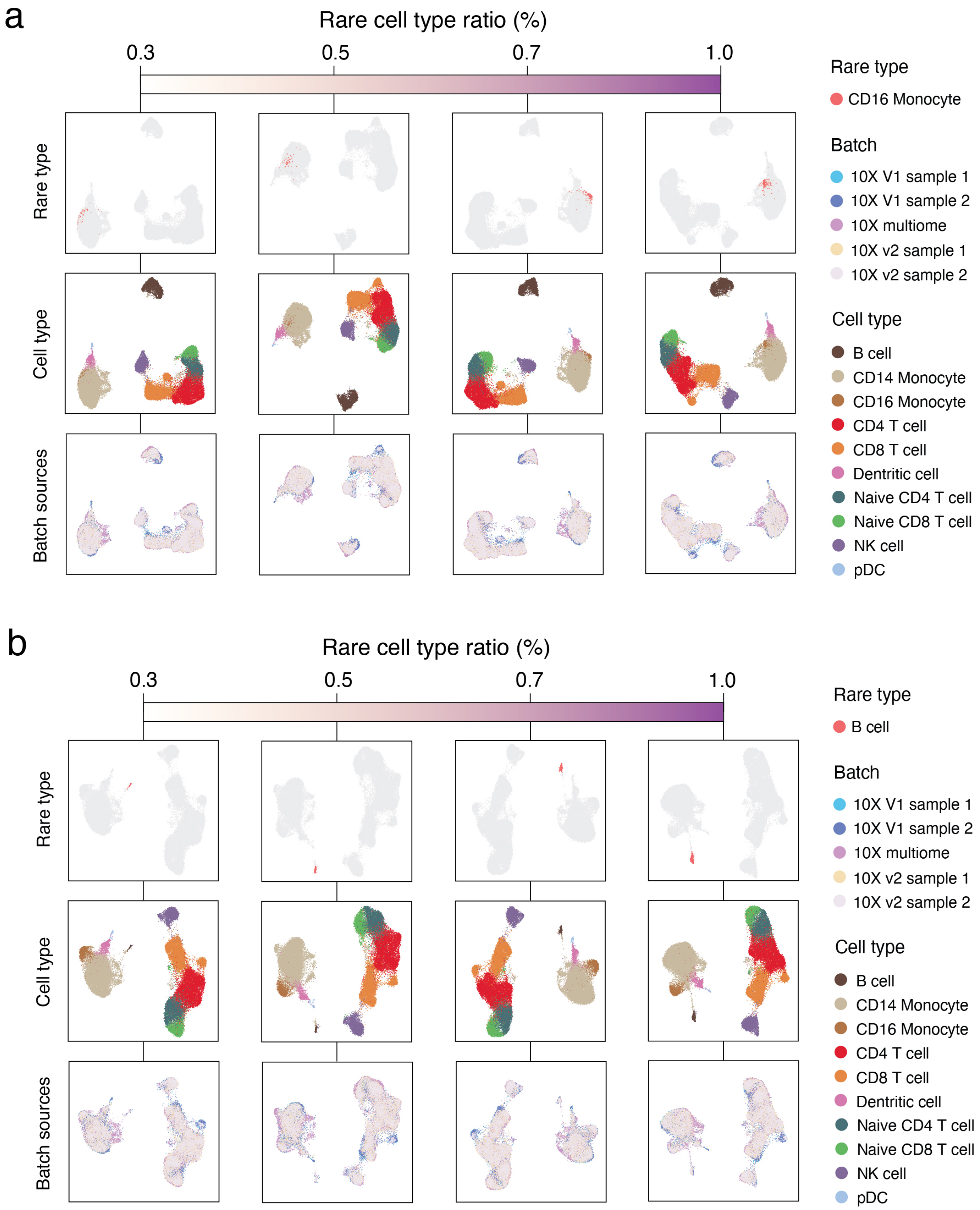

**Supplementary Fig. 9. a.** Benchmarking result for the experiment group "reference: 10x v2 and multiomics, query: 10x v1" (n=5 for 5 repeating experiments) **b.** Benchmarking result for the experiment group "reference: 10x v1 and v2, query: 10x multiomics" (n=5 for 5 repeating experiments)

##### 3.0.10 Supplementary Fig 10. UMAP visualization of the predicted labels against the ground-truth label on the query 2 experiment set.

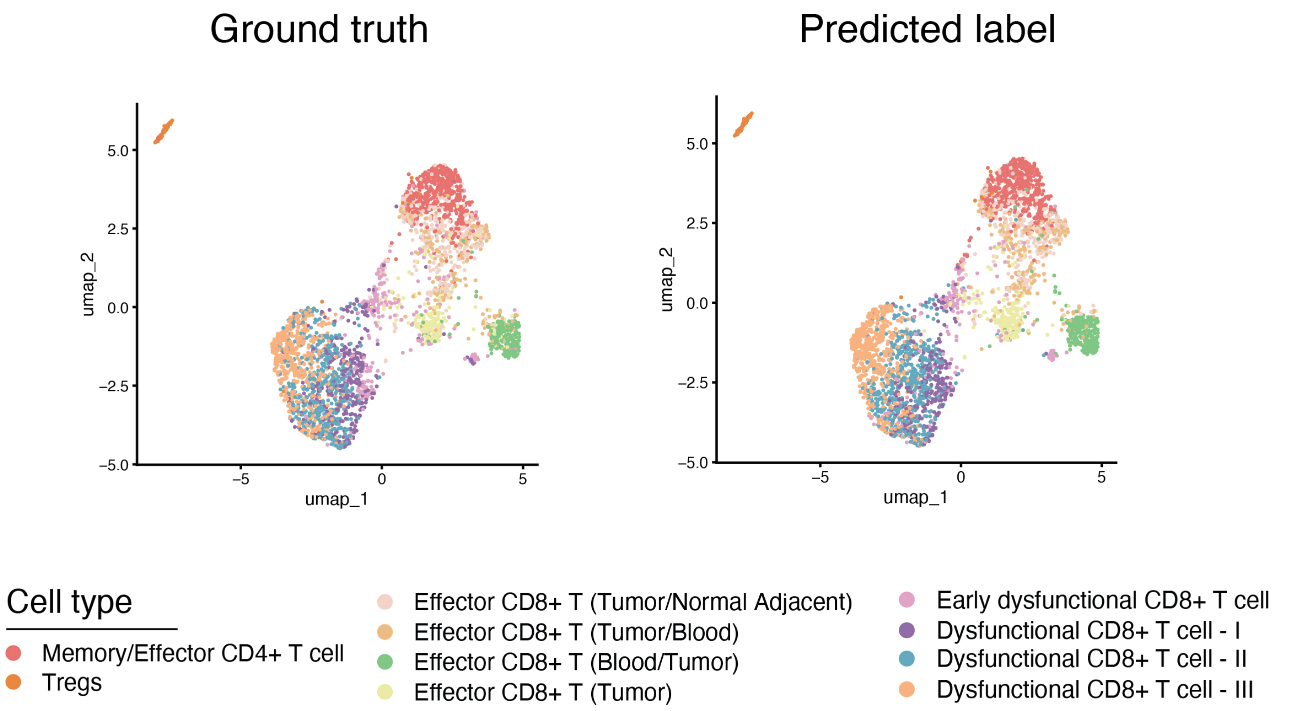

**Supplementary Fig. 10.** UMAP visualization of the predicted labels against the ground-truth label on the query 2 experiment set.

##### 3.0.11 Supplementary Fig 11. Boxplots of TPR and FDR benchmarking results with CD8 T cells as the out-of-reference (OOR) population.

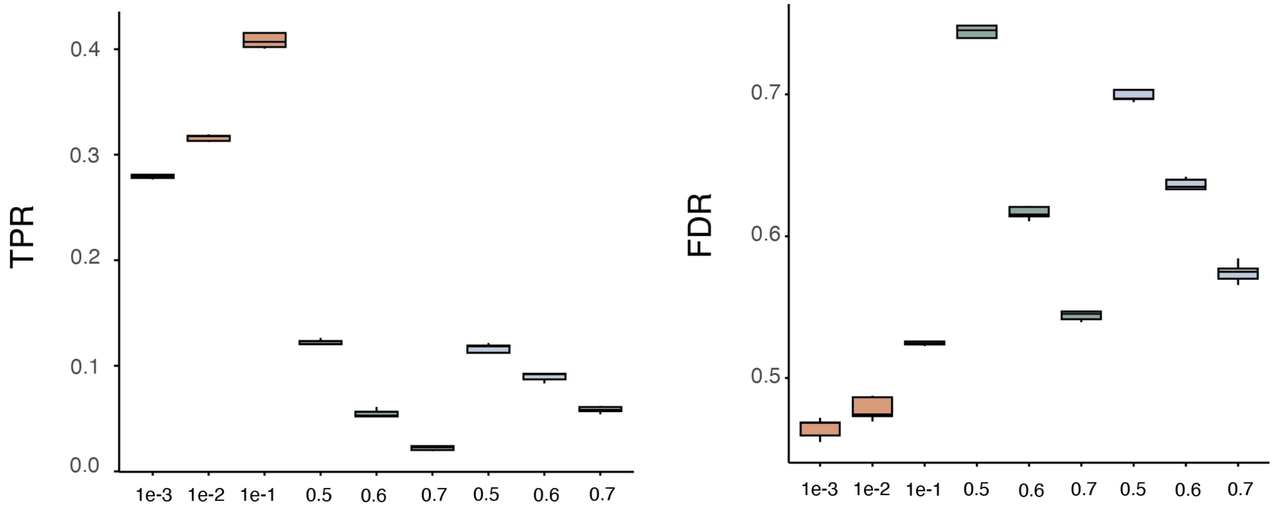

**Supplementary Fig. 11.** Boxplots of TPR and FDR benchmarking results with CD8 T cells as the out-of-reference (OOR) population. Orange - EpiPack, Green - kNN, Blue - SVM.

##### 3.0.12 Supplementary Fig 12. UMAP visualization of the gene score highlights in reference atlas embedding.

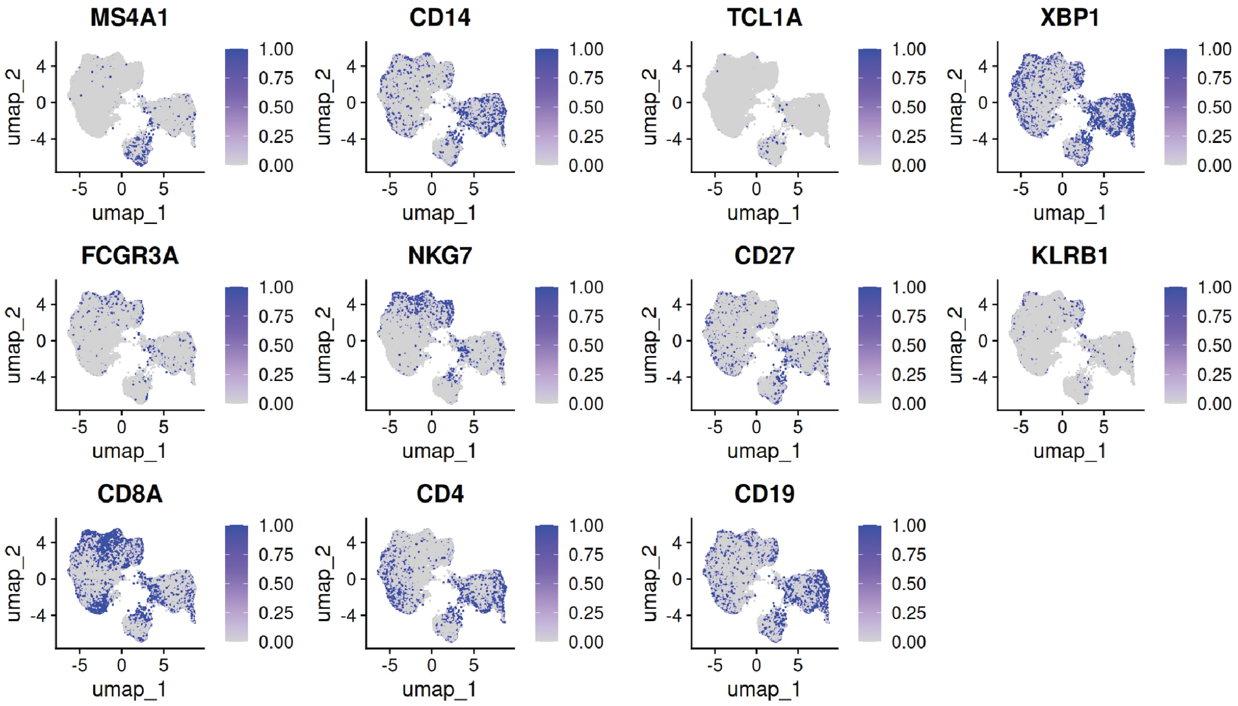

**Supplementary Fig. 12.** UMAP visualization of the gene score highlights in reference atlas embedding to confirm cell type annotation result.

##### 3.0.13 Supplementary Fig 13. UMAP visualization of the gene score highlights in the health-COVID joint embedding.

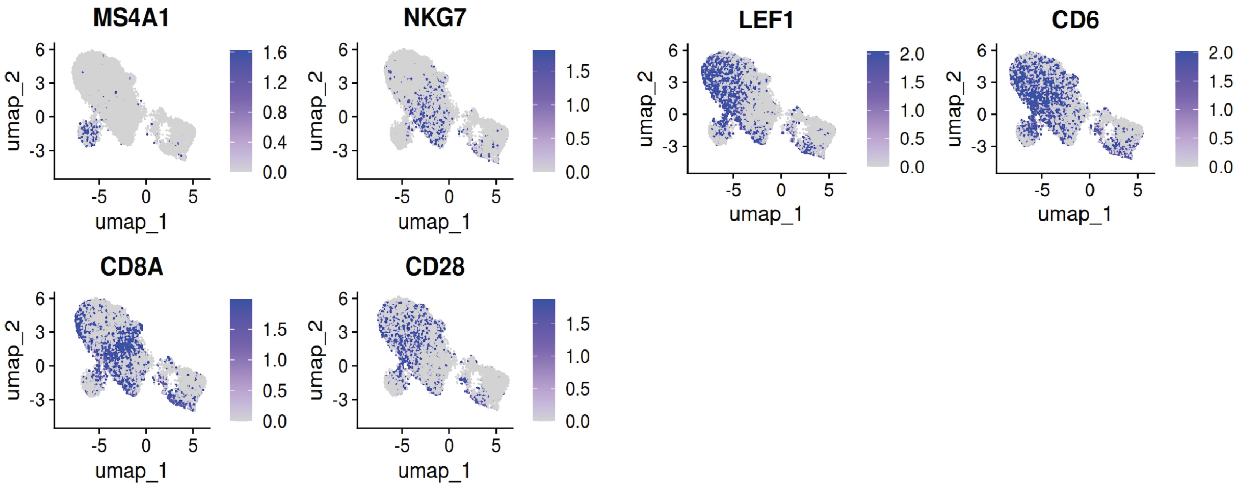

**Supplementary Fig. 13.** UMAP visualization of the gene score highlights in the health-COVID joint embedding to confirm cell label transfer result.

##### 3.0.14 Supplementary Fig 14. Differential peak analysis in the B cell between the OOR group and the health control cluster.

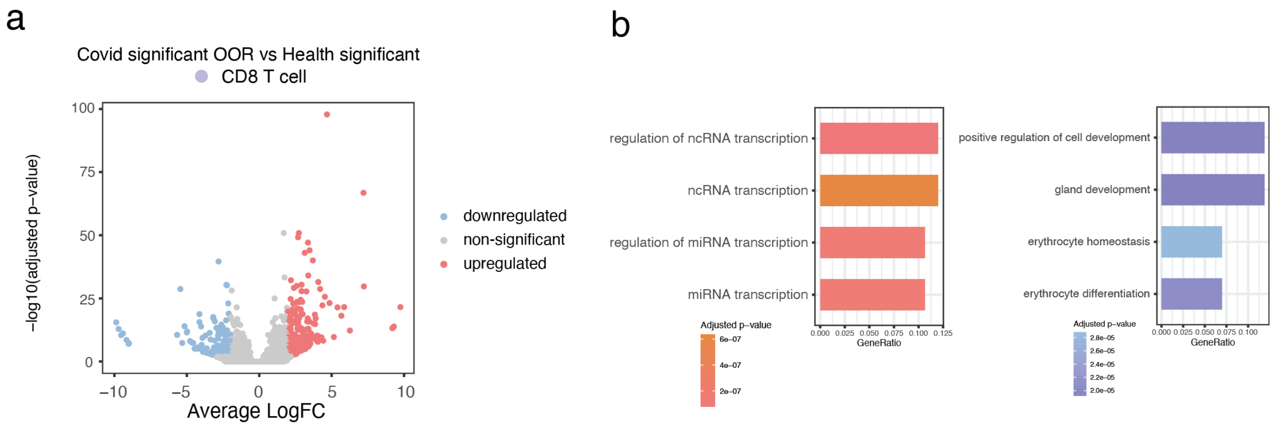

**Supplementary Fig. 14. a.** Volcano plot of differential peak analysis between healthy and COVID-associated B cells. **b.** Bar plots displaying the most significantly enriched GO biological processes associated with differential peaks in each cluster.
